# Microbial glutamate metabolism predicts intravenous cocaine self-administration in Diversity Outbred mice

**DOI:** 10.1101/2022.09.11.507297

**Authors:** Thi Dong Binh Tran, Hoan Nguyen, Erica Sodergren, Center for Systems Neurogenetics of Addiction, Price E Dickson, Susan N Wright, Vivek M Philip, George M. Weinstock, Elissa J. Chesler, Yanjiao Zhou, Jason A. Bubier

## Abstract

The gut microbiome is thought to play a critical role in the onset and development of psychiatric disorders, including depression and substance use disorder (SUD). To test the hypothesis that the microbiome affects addiction predisposing behaviors and cocaine intravenous self-administration (IVSA) and to identify specific microbes involved in the relationship, we performed 16S rRNA gene sequencing on feces from 228 diversity outbred mice. Twelve open field measures, two light-dark assay measures, one hole board and novelty place preference measure significantly differed between mice that acquired cocaine IVSA (ACQ) and those that failed to acquire IVSA (FACQ). We found that ACQ mice are more active and exploratory and display decreased fear than FACQ mice. The microbial abundances that differentiated ACQ from FACQ mice were an increased abundance of *Barnesiella, Ruminococcus*, and *Robinsoniella* and decreased *Clostridium IV* in ACQ mice. There was a sex-specific correlation between ACQ and microbial abundance, a reduced *Lactobacillus* abundance in ACQ male mice, and a decreased *Blautia* abundance in female ACQ mice. The abundance of *Robinsoniella* was correlated, and *Clostridium IV* inversely correlated with the number of doses of cocaine self-administered during acquisition. Functional analysis of the microbiome composition of a subset of mice suggested that gut-brain modules encoding glutamate metabolism genes are associated with the propensity to self-administer cocaine. These findings establish associations between the microbiome composition and glutamate metabolic potential and the ability to acquire cocaine IVSA thus indicating the potential translational impact of targeting the gut microbiome or microbial metabolites for treatment of SUD.

**HIGHLIGHTS:**

- Correlational analysis of novelty behaviors to IVSA acquisition shows that mice that acquire cocaine IVSA are more active and exploratory and have decreased fear than those that failed-to-acquire IVSA.
- The gut microbiome profiling of 228 diversity outbred mice indicates the relative abundances of *Barnesiella, Ruminococcus, Robinsoniella* and *Clostridium* IV are associated with the ability to self-administer cocaine.
- Associations between the gut microbiome and IVSA acquisition are sex-specific. Decreased relative abundances of *Lactobacillus and Blautia* are associated with IVSA in male and female mice, respectively.
- The relative abundances of *Robinsoniella* and *Clostridium* IV were correlated with the number of infusions of self-administered cocaine.
- Functional potential analysis of the gut microbiome supports a role for microbiomes encoding glutamate metabolism in the ability to self-administer cocaine.

## 1. Introduction

Substance use disorder (SUD) is a chronic brain disease characterized by compulsive drug seeking and use, despite negative consequences ^1^. While candidate genes and genome-wide association study (GWAS) yielded important insight into the genetic variation influencing addiction, little is known about the impact of the gut microbiome. There is growing awareness in the medical community that an imbalance of the gut microbiome (dysbiosis) is associated with various local and systemic diseases ^2^, including psychiatric disorders^3–5^. Indeed, compared to people who do not use cocaine, people who use cocaine have an altered gut microbiome.^6^ Furthermore, the gut microbiome can affect behavioral response to cocaine in mice ^7^. Mice treated with non-absorbable antibiotics exhibited enhanced sensitivity to cocaine reward and enhanced sensitivity to the locomotor sensitizing effects of repeated cocaine administration. Anxiety and depression are common comorbidities of SUD ^8^. The gut microbiome can induce anxiety and depressive-like behavior in mice, with the exact underlying mechanism remaining elusive. Metabolites, specifically short-chain fatty acids (SCFA) produced by bacterial fermentation of non-digestible carbohydrates, are important mediators in gut-brain crosstalk. 50% of the body’s dopamine and upwards of 90% of the body’s serotonin are produced in the intestinal tract ^9^. Although these two addiction-related neurotransmitters do not directly cross the blood-brain barrier (BBB), they may affect the brain’s reward centers through immune signaling or direct interaction with vagus nerves, resulting in neuroinflammation^10,11^. Administration of probiotic *Lactobacillus plantarum* PS128 increased levels of serotonin and dopamine in the striatum and resulted in changes in anxiety-like behaviors in mice^12^. The role of *Lactobacillus reuteri* in social behavioral defects was differentiated from the *Cntnap2-* dependent control of hyperactivity in mice^13^. Subsequent metabolomic studies of *Lactobacillus reuteri* inoculation showed biopterin rescues social interaction-induced synaptic transmission (as does cocaine) through stimulation of the vagus nerve^14^ and oxytocin release^15,16^. Together, these observations have resulted in translational advances, including clinical trials of microbiota-based interventions in autism spectrum disorder^17^. The potential of targeting the brain through the gut to address various neuropathology is becoming a reality^18^. However, the relevant microbes, small molecules and mechanisms of their action on behavior need to be identified. Mouse genetics provide a powerful and holistic approach for the detection of such targetable mechanisms of gut-behavior interaction.

The Collaborative Cross (CC)^19^ mouse panel represents a set of recombinant inbred strains of mice derived from a cross of eight parental inbred lines (five classic inbred strains and three wild-derived inbred strains) ^20–23^. These strains have been used to identify divergent responses to cocaine^24^. The Diversity Outbred (DO) mice were derived from CC mice outcrossed to maintain the distribution of alleles from across the eight parental strains in a unique combination in each mouse^23,25,26^. Each DO mouse is unique, and the population is heterogeneous, giving theoretically unlimited precision in genetic mapping to identify the source of variation in complex traits. The DO harbors variation in virtually all genes and pathways in the mouse, with 45 million variants. Using this population we can identify genes and microbes associated with variation in addiction related behaviors. In a small cohort of DO mice, we have previously shown using multivariate methods that various novelty behavior measures positively correlate with multiple cocaine intravenous self-administration behavior (IVSA) metrics^27^. IVSA of cocaine is considered the gold standard methodology to model volitional drug use in rodent studies ^28^. The propensity to self-administer addictive drugs has been repeatedly associated with predisposing traits including impulsivity and novelty-seeking behaviors^29–31^, allowing powerful assessment of the predisposing biological characteristics predictive of substance use disorders and related rodent phenotypes.

To identify microbes and microbial metabolites that are associated with the propensity to substance use disorders, we examine open field, light dark, holeboard and novelty place preference. Each of these novelty-related assays likely represents phenotypically distinct and genetically independent constructs^32^. After novelty-related testing, gut microbes were sampled before performing cocaine IVSA testing. We compared the microbiomes of mice that acquired IVSA (ACQ) to those that failed to acquire IVSA (FACQ) and related those microbes to the novelty behaviors. Finally, we explored the pathways that are predicted to differ between these groups using PICRUSt and Gut-Brain Module analysis of mWGS data.

## 2. Methods and Materials

### 2.1 Animals

In total, a large DO population of 1,065 mice were evaluated on a set of behavioral phenotyping assays. For the microbiome analysis, 228 J:DO (The Jackson Laboratory JR009376) from both sexes (M=121, F=107), spanning generations G21, G22 and G23, were purchased from the Jackson Laboratory in room G200, a high barrier Pathogen & Opportunistic-Free Animal Room (Health report https://www.jax.org/-/media/jaxweb/health-reports/g200.pdf?la=en&hash=7AD522E82FA7C6D614A11EFB82547476157F00E1) and transferred at wean to an intermediate barrier specific pathogen-free room (https://www.jax.org/-/media/jaxweb/health-reports/g3b.pdf?la=en&hash=914216EE4F44ADC1585F1EF219CC7F631F881773). Same-sex siblings were co-housed with a combined body weight of up to 15 g per side of the duplex cage at 3-5 weeks. Mice were individually housed starting at six weeks of age with a minimum of 5 days before testing. Mice were housed under (12:12) light-dark cycle and allowed *ad libitum* access to the standard rodent chow [sterilized NIH31 5K52 6% fat chow (LabDiet/PMI Nutrition, St. Louis)] and acidified water (pH 2.5–3.0) supplemented with vitamin K. Mice were housed in duplex individually vented Thoren cage #3, cages with pine-shaving bedding (Hancock Lumber) and environmental enrichment consisting of a nestlet and Shepard’s shack. The mice were identified by ear notching at 3-4 weeks of age and moved between cages and testing arenas using metal forceps. Clean housing was provided through a cage change once a week. If this fell on a testing day, then a clean cage was prepared, and post-test, the mice were housed in their new clean cage. The Jackson Laboratory follows husbandry practices in accordance with the American Association for the Accreditation of Laboratory Animal Care (AAALAC), and all work was done with the approval of our Institutional Animal Care and Use Committee (Approval #10007).

### 2.2 Phenotyping

At three to six months, mice were phenotyped using the open-field, light-dark, holeboard, and novelty place preference assays daily from Monday to Thursday for four separate times. After novelty testing, mice underwent jugular catheter implantation surgery to prepare for cocaine intravenous self-administration (IVSA). Complete phenotyping details can be found at https://mpdpreview.jax.org/projects/CSNA03/protocol

#### 2.2.1 Open Field Apparatus

A square-shaped, clear polycarbonate arena (Med-Associates #MED-OFAS-515U) with dimensions 17.5 inches length x 17.5 inches width x 10.0 inches height (44.5 cm x 44.5 cm x 25.4 cm) was used for the open field arena. External to the arena’s perimeter at the floor level, on the left and right sides, were a pair of horizontal infrared photobeam sensors (16 × 16 beam array). An additional pair of infrared photobeam sensors raised 3 inches from the arena floor (16 × 16 array) were situated at the front and rear external sides of the arena and used to capture vertical activity. Each arena was placed within a sound attenuated, ventilated cabinet with interior dimensions: of 26”W x 20”H x 22”D (Med Associates, #MED-OFA-017). Each cabinet contained two incandescent lights, each affixed in the upper rear two corners of the cabinet at the height of approximately 18.5 inches from the center of the arena floor, providing illumination of 60±10 lux when measured in the center of the arena floor. Data was collected using Activity Monitor software:7.0.5.10 SOF-812 (Med Associates, Inc; RRID:SCR_014296). For all behavioral assays, mice were transported from the housing room to the procedure room on a wheeled rack and left undisturbed to acclimate to the anteroom adjacent to the procedure room for a minimum of 30 minutes. Before the first mouse was placed into any arena and between subjects, the chamber was thoroughly sanitized with 70% ETOH solution (in water), and the box was wiped dry with clean paper towels. After all testing for the day, the subjects were returned to the housing room, and the arenas were sanitized with Virkon followed by 70% ETOH to remove any Virkon residue.

##### 2.2.1.1 Open field activity

At the start of each testing session, mice were individually placed in the center of the arena, facing the rear of the chamber. The lid was placed atop the arena, and the chamber door was closed. The tracking software detects the mouse in the arena and starts automatically. The tracking software automatically ends the tracking for the subject 60 minutes after the mouse was initially detected by the software. Numerous metrics were recorded for each mouse see data definition **Sup Table 1**.

##### 2.2.1.2 Light-dark

A black polycarbonate insert that does not interrupt infrared beams (Med Associates) was placed into the back half of the open field arena, with the entry door opening toward the arena’s front center, thereby reducing the light levels within the dark side of the box. At the start of each testing session, mice were individually placed in the center of the arena, on the light side, facing the rear of the chamber (facing the dark side). The lid was placed atop the arena, and the chamber door was closed. The tracking software detected the mouse in the arena and started automatically. The tracking software automatically ends the tracking for the subject 20 minutes after the mouse was initially detected by the software. Numerous metrics were recorded for each mouse see data definition **Sup Table 1**.

##### 2.2.1.3 Holeboard

The holeboard floor (Med Associates) was inserted into the open field arena. The holeboard floor consists of two components: the metal floor, which the mice were in contact with, was comprised of 16 evenly spaced holes, and the under plate contains 16 holes matching those on the metal plate with wire mesh for baiting the holes. At the start of each testing session, subjects were individually placed in the center of the arena, facing the rear of the chamber. The lid was placed atop the arena, and the chamber door was closed. The tracking software detected the mouse in the arena and started automatically. The tracking software automatically ended the tracking for the subject 20 minutes after the mouse was initially detected by the software. Numerous metrics were recorded for each mouse see data definition **Sup Table 1**.

#### 2.2.2 Novelty place preference

The Novelty Place Preference apparatus was a rectangular shaped, three-chambered arena (acrylic or polycarbonate materials) with dimensions (white/black chambers - 5 in length, 6.5 in width, 5 in height, grey center chamber – 5 in length, 3.5 in width, 5 in height) with automatic doors between the three sections, and a clear, aerated lid (Med Associates, Inc.) The bottom of the arena contained pairs of horizontal infrared photobeam sensors at the floor level, which were not visible to the mouse. Each arena was placed within a sound attenuated, ventilated cabinet with dimensions 25 in width x 26 in depth x 21 in height (Med Associates, Inc.) Before the start of testing, upon transport to the procedure room, subjects were briefly handled and assessed for welfare concerns that may have resulted in exclusion from testing (e.g., fight wounds or bite marks). They were then left undisturbed to acclimate to the procedure room environment for a minimum of 60 min before testing began. The mouse was placed in the center grey chamber, and all hinged ceiling doors were closed. Once the software detected the placement of the mouse, the ceiling light in the chamber was turned off, and a 5-minute acclimation timer was started. After the 5-minute acclimation period, the motorized guillotine doors leading to the preassigned exposure chamber (black or white) opened, and the ceiling lights turned on. After 20 minutes, the exposure session was complete; the guillotine door closes. Immediately upon the conclusion of the exposure program, the mouse was placed back into the center compartment. If a mouse was in the center chamber after the exposure program as above, the mouse was handled and removed from the center chamber and then replaced in the center chamber to mirror the handling of all subjects between exposure and test programs. Upon all subjects being returned to the program immediately detected the mice, the ceiling light in the chamber turned off, and a 5-minute acclimation timer started. After the 5-minute acclimation period, *both* motorized guillotine doors leading to the black and white chambers opened, and the ceiling lights turned on. A 10-minute testing timer begin. After 10 minutes, the session was complete, and the doors closed. Only one metric was recorded: the % movement on the novel side.

### 2.3 Fecal Collection

After the four days of behavioral testing, fecal pellets were collected from the mice before surgery on day five. Mice were placed in a clean wean pen for five minutes. Any fecal pellets deposited were immediately collected, placed in Eppendorf tubes, and stored at −80 °C until DNA extraction.

### 2.4 Cocaine Intravenous self-administration (IVSA)

Mice were tested following a modification of the protocol described in Dickson et al. ^27^. Mice > 12 weeks of age underwent indwelling jugular vein catheterization under oxygen/isoflurane anesthesia by The Jackson Laboratory surgical services. The catheter (Instech, C20PU-MJV, Plymouth Meeting, PA) was routed through the stainless coupler of a mesh button. The mesh button was sutured in place subcutaneously. The port was flushed with saline and filled with 10 ul of lock solution (Lumen lock solution; 500 IU heparin/ml lock solution). The catheter was capped with a protective aluminum cap (Instech., PHM-VAB95CAP). The mouse was observed for at least three days after surgery to ensure the incision was healing properly. If mice exhibited pain signs, buprenorphine (0.05mg/kg SQ) was administered every 4-5 hours or carprofen (5 mg/kg SQ) was administered every 24 hours. Mice were provided a minimum ten-day post-operative recovery period in their home cage before testing. Mice were transported in their home cages from the housing room to the procedure room on a wheeled transport rack. Before the start of testing, upon transport to the procedure room, mice were weighed (once a week) and briefly handled to be assessed for any welfare concerns that may result in exclusion from testing (e.g., wounds or looking sickly). IVSA data were collected using Med Associates operant conditioning chambers (307W), fitted with two retractable levers on the front wall flanking the right and left sides of the center panel food hopper. Directly over each lever (~ 2-3 inches above) were red stimulus lights. A house light (ENV-315W) was centrally mounted on the rear wall. A modified Plexiglas floor, fabricated at the Jackson Laboratory, was fitted to cover the metal floor grids. The chambers were within sound attenuating cubicles (ENV-022MD). A 25-gauge single-channel stainless steel swivel was mounted to a counterbalanced lever arm attached to the outside of the chamber. Tubing was used to connect a syringe mounted on the infusion pump to the swivel and to connect the swivel to a vascular access harness. Operant conditioning chambers were controlled by a Med Associates control unit using MED-PC IV software (RRID:SCR_012156).

Mice began cocaine IVSA testing on a fixed-ratio 1 (FR1) schedule at a dose of 1.0 mg/kg/infusion of cocaine hydrochloride (NIDA Drug Supply). Infusions from the active lever were at 8.85 ul/sec, with 1 sec per 10g mouse weight. Each IVSA session was two hours each day. After each day of testing, Baytril (enrofloxacin) was administered by catheter to the mice at 22.7 mg/kg, followed by a 20 ul injection of Heparin lock solution (100 U/ml heparin/saline). Acquisition criteria were met when mice had 5 (could be non-consecutive) sessions with ≥10 infusions. Mice were excluded from the study if they did not acquire within 18 days of starting. A mouse may be excluded earlier if it can’t meet the acquisition criteria within 18 days. The mouse reached stabilization when the number of infusions didn’t vary more than 20% for the last two consecutive sessions. Following stabilization, the mice were tested on 1.0 mg/kg through a dose-response curve. During this testing phase, the mice were assessed for self-administration response across a 4-dose cocaine dose-response curve (DRC). Doses were presented in the following order: 1.0, 0.32, 0.1, 0.032 mg/kg/infusion. Mice were tested on consecutive days on the same dose until stabilization criteria were met or a max of 5 sessions, then moved on to the next dose. Following the DRC, mice were evaluated for extinction, defined as a reduction in lever presses on the active lever that does not result in a cocaine infusion reward.

During extinction sessions, subjects were connected to saline-filled infusion tubing; however, the infusion pump was not activated. During extinction sessions, the house light was continuously illuminated, stimulus lights were never illuminated, and lever presses have no programmed consequences. Mice were tested for 3-9 days, depending on whether the extinction criteria are met. The number of active lever presses on the first day of the extinction session was the reference baseline for establishing extinction. The criteria for extinction were defined as follows: a) When active lever presses are reduced to <50% of the established baseline from the initial extinction session; and b) the variance of the final two days is within 20% of the established baseline from the initial extinction session; c) if active lever presses are <10 on or after day 3. Subjects not reaching extinction criteria within nine days were advanced to the reinstatement stage. Following extinction, responding on the active and inactive levers was examined for two daily two-hour sessions. During reinstatement sessions, as in the acquisition and DRC, drug-paired stimuli, including the sound of the infusion pump, were presented; however, there was no infusion into the mouse. Specifically, mice were connected to infusion tubing, which was connected to a sterile-saline-filled syringe, but the syringe was not in the pump. Thus, mice do not receive drug infusions. Initiation of lever pressing was due to conditioned stimuli. Numerous metrics were recorded; see data definition **Supp Table 1**. Not all mice that entered the study completed it. Mice may fail to survive the surgery, have a non-patent catheter, a systemic infection, a local infection at the catheter port, cocaine overdose, or air embolism. These mice were excluded from the study.

### 2.5 16S rRNA gene sequencing, data processing and microbiome functional prediction

Shoreline Complete V1V3 kit (Shoreline Biome, cat #SCV13) https://www.shorelinebiome.com/product/v1-v3-16s-amplicon-kit/) was used per manufacturer’s instructions for DNA extraction, library preparation and sequencing. Briefly, ~5mg of mouse fecal pellet was lysed with a combination of heat, pH, and cell wall disruptors in a single step. DNA was recovered using the supplied magnetic beads, and an aliquot from each sample was transferred to a well in the provided PCR plate containing sample barcode/Illumina sequencing primers. 2x PCR mix from the kit was added, and PCR was performed according to the manufacturer’s instructions. After PCR, samples were pooled, purified using a MinElute PCR Purification Kit (Qiagen, cat# 28004) and diluted for sequencing on the Illumina MiSeq platform generating 2×300 paired reads.

Sequencing reads were processed by removing the sequences with low quality (average qual <35) and ambiguous codons (N’s) using the in-house pipeline. Paired amplicon reads are assembled using Flash 2.0. Chimeric amplicons were removed using UChime software (RRID:SCR_008057)^33^. Our automatic pipeline used the processed reads for operational taxonomic unit (OTU) generation based on a sequence similarity threshold of 97%. Each OTU was classified from phylum to genus level using the most updated Ribosomal Database Project (RDP) 2016 classifier and training set (RRID:SCR_006633). A taxonomic abundance table was generated with each row as bacterial taxonomic classification, each column as sample ID and each field with taxonomic abundance. The abundance of a given taxon in a sample was present as relative abundance (the read counts from a given taxon divided by total reads in the sample). We also used PICRUSt2 (RRID:SCR_022647)^34^ to predict the functional profiling of the bacterial communities by ancestral state reconstruction using 16S rRNA gene sequences. Following the protocol described by Valles-Colomer^5^, gut-brain module analysis was performed on the PICRUST2 results. Multiple testing correction of the Gut-Brain Modules was performed using the qvalue package (RRID:SCR_001073).

### 2.6 mWGS sequencing, Data Processing and Metagenomics Species Detection

Libraries are constructed with an average insert size of 500 bases and then sequenced on the HiSeq2500 instrument producing 150 base read pairs from each fragment yielding ~3 million read pairs/sample. Following mWGS sequencing, sequence data was run through a quality control pipeline to remove poor-quality reads and sequencing artifacts. Sequencing adapters were first removed using TRIMMOMATIC (RRID:SCR_011848)^35^. Next, exact duplicates, low quality, low complexity reads and mouse DNA contamination were removed using GATK-Pathseq (RRID:SCR_005203)^36^ pipeline with a k-mer based approach. For optimization of mouse DNA decontamination, we have built a new GATK-Pathseq 31-mer database by concatenating the following collection of DNA sequences: MM10_GRCm38 reference; sixteen diverse laboratory mouse reference genomes define strain-specific haplotypes and novel functional loci; NCBI UniVec clone vector sequences; repetitive element sequences from RepBase23.02 database; and mouse gencode transcripts databases (v25). The final clean reads were used for taxonomic classification and metabolic function analysis for further downstream analysis.

An optimized GATK-Pathseq classification pipeline is time efficient and robust solution for taxonomic classification at the species level. This pipeline used BWA-MEM alignment (minimum 50 bp length at 95% identity). It mapped the final clean reads to the latest updated reference of microbial genomes built by concatenating RefSeq Release 99 (March 2nd, 2020) nucleotide FASTA sequence files of bacteria, viruses, archaea, fungi, and protozoa. Gatk-Pathseq All read counts of microbial species of all kingdoms are used for species abundance analysis.

### 2.7 Differential abundance analysis of microbial genes and metabolic pathways

The KEGG ortholog (KO) profiling was performed by HumanN2 (RRID:SCR_016280)^37^. Using DESeq2 package (RRID:SCR_015687)^38^ dedicated to performing comparative metagenomics, the inference of abundance of genes and pathways was obtained and visualized using volcano plot. Because of the potential high false positive rate of DESeq^39^, we plotted raw and relative abundance to inspect the results. Following the protocol described by Valles-Colomer^5^, gut-brain module analysis was performed on the mWGS results.

Multiple testing correction of the Gut-Brain Modules was performed using the qvalue package (RRID:SCR_001073).

### 2.8 Statistical Analysis

High-throughput behavioral phenotyping data is known to be affected by unwanted sources of variation such as the date of testing, seasonal effects and the effect of the tester. Before using this behavioral phenotyping data, we assessed and corrected for these. First, we performed an outlier analysis removing any phenotypes greater than 3 SD of the mean. Then we tested to determine if the tester or date of the test had significant effects, p<0.05. If those variables were significant, they were added to a model, the residuals of which were then tested for normality. If those residuals were normally distributed, they would be used for downstream analysis. If the residuals were not normally distributed, they were rank Z transformed. This process was performed on a large dataset of over 3,000 DO mice. A subset of which were used in IVSA and a subset of those for this microbiome analysis.

Microbial community analysis was performed by R version 3.5.1 (RRID:SCR_001905). Principal coordinate analysis (PCoA) plots, boxplots and heatmaps were generated for graphical visualization using Phyloseq, ggplot2 (RRID:SCR_014601)^45^ and ComplexHeatmap (RRID:SCR_017270)^46^ packages. Richness was calculated as the number of OTUs present in each sample. The Shannon Diversity Index combined species richness, and the evenness was computed as ∑p_i_*ln(p_i_), where pi presents the proportional abundance of species. The non-parametric Wilcoxon or Kruskal-Wallis rank sum-tests were used for differential diversity or abundance between two or more groups and corrected for multiple comparisons by the Benjamini-Hochberg procedure. Beta diversity was analyzed at the OTU level using the Bray-Curtis distance for community abundance and the Jaccard distance for community presence/absence.

The among-group differences were determined using the permutational multivariate analysis of variance by the distance matrices (ADONIS). These tests compare the intragroup distances to the intergroup distances in a permutation scheme and then calculate a p-value. These functions are implemented in the Vegan package (RRID:SCR_011950)^41^. For all permutation tests, we used 10,000 permutations.

Association between taxa and behavioral variables was assessed by fitting the Generalized Linear Model-GLM (glm R function). This approach raises additionally interesting associations between bacteria and behavior related to Age and Sex. The residuals were not checked for normality rather the GLM regression coefficients were standardized using lm.beta R function. The Benjamini-Hochberg procedure (FDR) was used to correct for multiple testing of taxa and behavioral test associations, with significance defined as FDR < 0.1.

We also used multi-task learning for feature selection by fitting a GLM via penalized maximum likelihood model (LASSO-Least Absolute Shrinkage and Selection Operator) (glmnet R package RRID:SCR_015505)^42^. The GLM allows us to predict the association of a single microbe with a novelty test regarding sex and age covariates. The p-value will be adjusted for multiple testing. By contrast, LASSO assesses the influences of all dependent variables on all response variables, and no assumption was made. In our analysis. We pooled all 42 known genera as dependent variables against 35 novelty behaviors as response variables. Using LASSO, the estimates of the regression algorithm are shrunk toward zero by adding a penalized term in the loss function. Relevant genera with nonzero coefficients were selected as informative variables associated with all behavior variables from the model. Thus, LASSO model represent an alternative approach of GLM to identify bacteria that associated with novelty behavior results.

### 2.9 Network analysis

Network analysis was performed using SParse InversE Covariance Estimation for Ecological Association Inference (SPIEC-EASI) (RRID:SCR_022646), which combines data transformation developed for compositional data and a graphical model inference framework assuming the underlying ecological association network^43^. The network was built on filtered ACQ and FACQ microbial data (filtered 50% OTU prevalence) of DO mice. All novelty behaviors were integrated as equal to each OTU.

## 3. Results

### 3.1 Differences in novelty-related behavior between mice that acquire self-administration of cocaine and those that failed-to-acquire self-administration of cocaine

Each of the novelty-behavior phenotyping paradigms results in numerous measurements. Therefore, we selected the most non-redundant yet still highly correlated measures of novelty; 24 open-field measures, six light-dark box measures, three hold board measures, and one novelty place preference measure to be correlated with the ability to acquire IVSA in a cohort of diversity outbred mice (n=228). For the IVSA assay, mice were categorized as having acquired cocaine self-administration (acquired (ACQ) n=128) if they reached the established criteria and were able to proceed to a dose-response curve, extinction and reinstatement or they were categorized as failed-to-acquire (failed-to-acquire (FACQ) n=100), in which case they never achieved the criterion of having acquired cocaine self-administration.

12 of the 24 open field measures were significantly different (p<0.05) between the ACQ and FACQ mice. The direction of these differences suggests that ACQ mice are more active and exploratory (e.g. total distance traveled W=7882, Z= −2.99 p=0.003, r=0.19) and less anxious (e.g. total time in the center W=7863, Z= −2.95, p=0.003, r=0.19) than FACQ mice ^47,48^ (**Fig. 1A, Sup Table 2)**. For the light-dark assay, two measures, the percent resting time in the light (W=7711, Z=−2.65, p=0.008, r=.17) and the percent time in the light (W=7670, Z=−2.57, p=0.01, r=0.17), were significantly different between ACQ and FACQ mice. Again, the direction of these differences suggests that ACQ mice are less fearful of the light side than FACQ mice^47^ **(Fig. 1B, Sup Table 2**.) Of the three holeboard measures, only the total number of entries was significantly different between ACQ and FACQ mice (W=7385.5, Z= −1.99, p<0.05, r=0.13), with ACQ mice having a greater number of hole pokes than FACQ mice. A high number of hole pokes are observed in mice given anxiolytics^48^ suggesting that the FACQ mice with fewer hole pokes are inherently more anxious (**Fig. 1C, Sup Table 2)**. The measure of novelty place preference, proportion of movement on the novel side, was significantly different (W=5335, Z=−2.15, p=0.03, r=0.14) between groups. In this case, the FACQ mice preferred the novel side over the familiar side, suggesting that ACQ mice had a more novelty-place aversion, which is associated with increased anxiety or anhedonia^49^ (**Fig. 1C, Sup Table 2)**.

**Figure 1.**
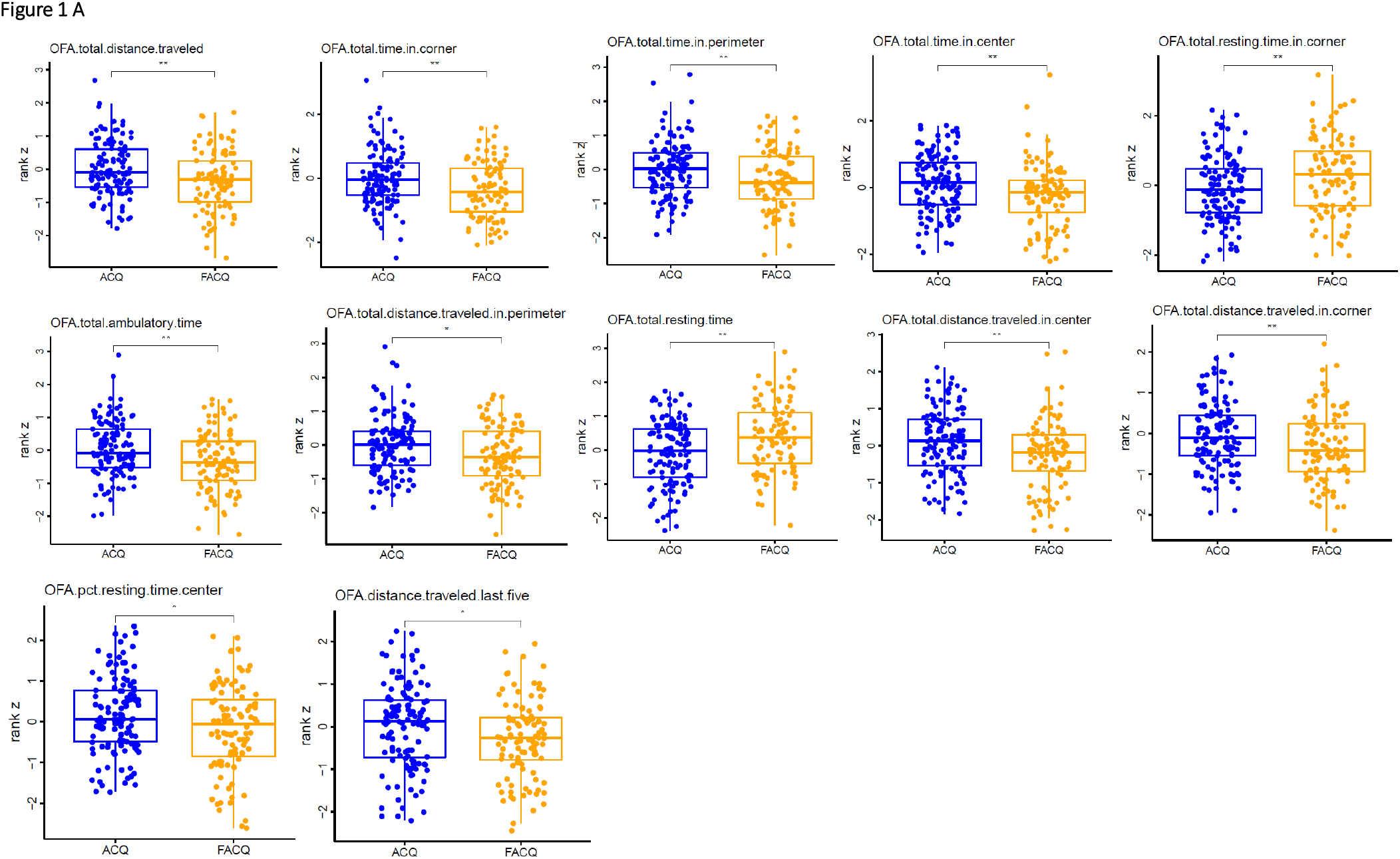
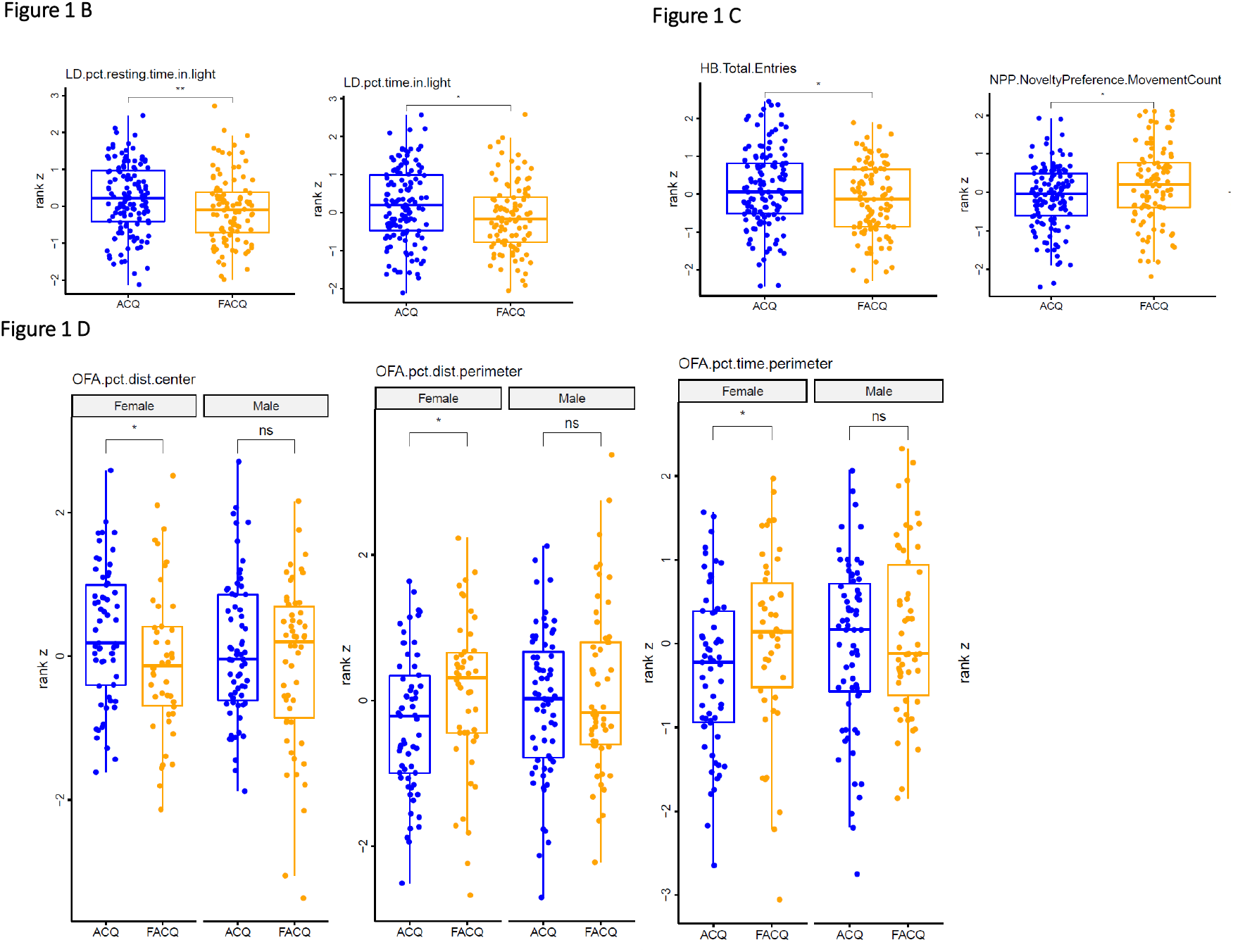
Novelty behaviors are associated with the acquisition of IVSA. **A)** The twelve open field measures differed between ACQ and FACQ. **B)** The two light-dark metrics that differed between ACQ and FACQ. **C)** The hole board total entries and the preference for a novel arena differed between ACQ and FACQ mice. **D)** Sex specificity novelty behaviors associated with the acquisition of IVSA. Three open field measures were significantly different in females, the percent distance and time in the perimeter and the percent distance time in the center. Outliers have been removed from the phenotype data, and the values rank z transformed. *p<0.05, **p<0.01.

### 3.2 Sex differences in novelty behavior between ACQ mice and FACQ

Mice of both sexes were tested, and when separated by sex, other behavioral differences were detected as sex-specific and not identified when pooled. Three open field measures differ between ACQ and FACQ considering sex. Female FACQ mice in spent significantly less time in the center than female in ACQ (W=1055, Z=−2.18, p=0.03, r=0.21) and a greater distance in the perimeter (W=1727, Z=−2.04, p=0.04, r=0.19), as well as percentage of time in the perimeter (W=1049, Z=−2.22, p=0.03, r=.21) (**Fig. 1D, Sup Table 2)**. The behaviors suggest that female ACQ mice are more anxious than female FACQ mice. We did not observed any male specific associations.

### 3.3 Microbial composition of the DO mice

The sampling depth average over 228 fecal samples was about 57,000 reads per sample. By using a cutoff of 25% of prevalence, we finally had 347 OTUs. The rarefaction curve shows highly consistent in terms of read depth related to species richness (data not shown). Firmicutes and Bacteroides were two major phyla of fecal pellets from the DO mice, with 58% and 34% relative abundance, respectively. *Firmicutes* to *Bacteroides* ratio was −4.9 (0.1-61.8). *Actinobacteria* is the third the most abundant phylum with about 7% of relative abundance, and six dominant genera presented 71% of microbial composition in the colon, such as *unclassified_Lachnospiraceae* (16.8%), *unclassified_Porphyromonadaceae* (15.6%), *Lactobacillus* (13%), unclassified*_Bacteroidales* (12.1%), *Eisenbergiella* (8%) and *Barnelsiella* (6.5) (**Sup Fig 1)**. The cohort had an alpha diversity of 236 (111-320) for richness.

### 3.4 Microbial composition of the ACQ vs. FACQ groups of DO mice

There were no significant differences in the richness and Shannon diversity between two IVSA groups when separated by sex or combined. Jaccard distance for community presence/absence showed no difference between groups (data not shown). The microbial composition and abundance of DO mice stratified by sex and groups were illustrated in Fig 2A (**Fig 2A**). In general, the bacterial community was highly dissimilar and reflected a stronger inter-individual variability, which is expected in an outbred mouse population PC1 accounted for 19.2% of the variance and PC2 13.7% of the variance of total microbiome community (**Fig 2B**). PERMANOVA analysis showed overall microbiome community structure was significantly different between two IVSA groups (p=0.035). Variance analysis revealed that IVSA groups accounted for only 0.8% (p=0.035) of microbiome variation, smaller than the effects sex (R2=1.1%, p=0.0018) and age (R2=2.2%, p=9.999e-05). At the phylum level, the relative abundance of *Firmicutes* that contain many butyrate-producing bacteria was lower in the ACQ group (56%) than in the FACQ group (60.8%). On the other hand, the relative abundance of Bacteroidetes was inversely higher in the ACQ group (36.4%) than in the FACQ group (30.9%).

**Figure 2.**
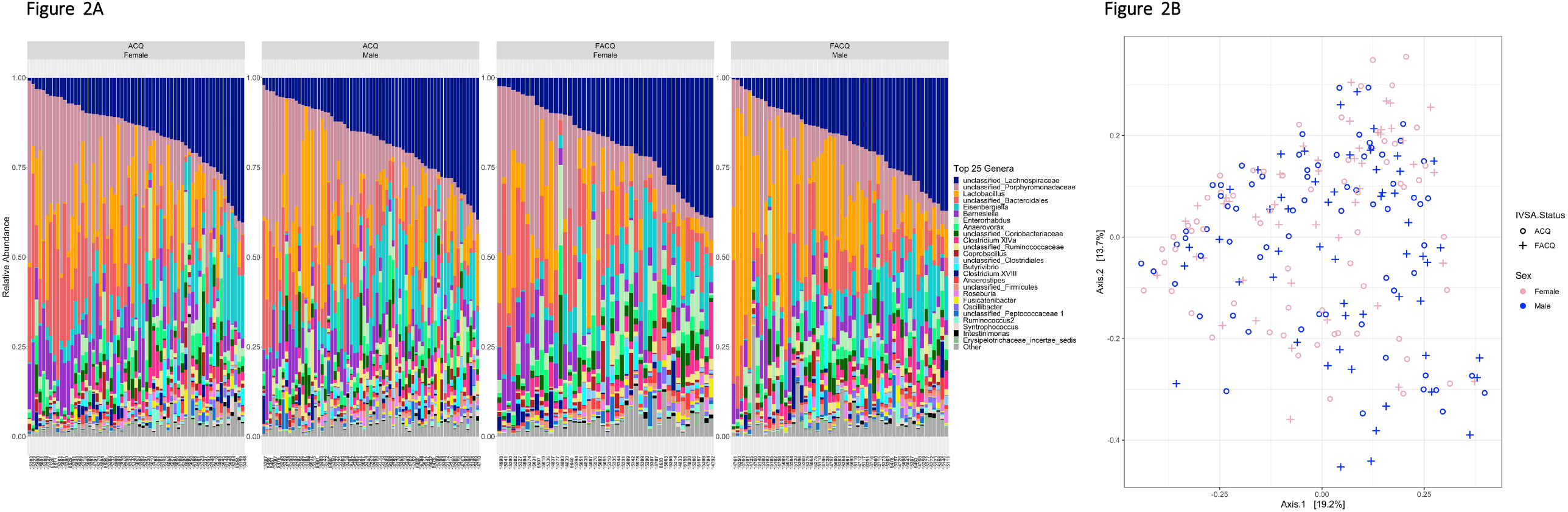
Microbial composition of the ACQ vs. FACQ groups of DO mice. **A)** The percent abundance of the 25 most abundant genera in female and male mice ACQ and FACQ mice. B) Principal components analysis of bacterial beta diversity at OTU level using Bray-Curtis for 16S microbiome data showing the ACQ, FACQ, and male and female groupings.

Four bacterial genera were significantly different between the two groups including the dominant taxon *Barnesiella* (W=7437, Z=−2.09, p= 0.035, r=0.138) and three other more rare microbes *Ruminococcus* (W=7801, Z=−2.15, p=0.004, r=0.19), *Robinsoniella* (W=7458, Z=−2.15, p= 0.03, r=0.14), and *Clostridium* IV (W=5321, Z=−2.18, p=0.03, r=0.14) (**Fig 3 A-D**). The abundance of *Blautia* was higher in female FACQ mice (W=1086.5, Z=−2.02, p=0.04, r=0.19), whereas *Lactobacillus* (W=1319, Z=−2.55, p=0.01, r=0.17) differed only in males where mice that FACQ had higher levels than ACQ mice (**Fig 3E-F**). One quantitative metric that all mice tested on IVSA had in common was the number of doses administered under the 1.0 mg/kg acquisition dose. Any mouse that failed to acquire would have less than ten doses administered. The doses administered across all mice and relative abundances of *Robinsoniella* (R=0.15) and *Clostridium IV* (R=−0.13) were significantly correlated (p<0.05) (**Fig. 3GH**, Sup Fig. 4).

**Figure 3.**
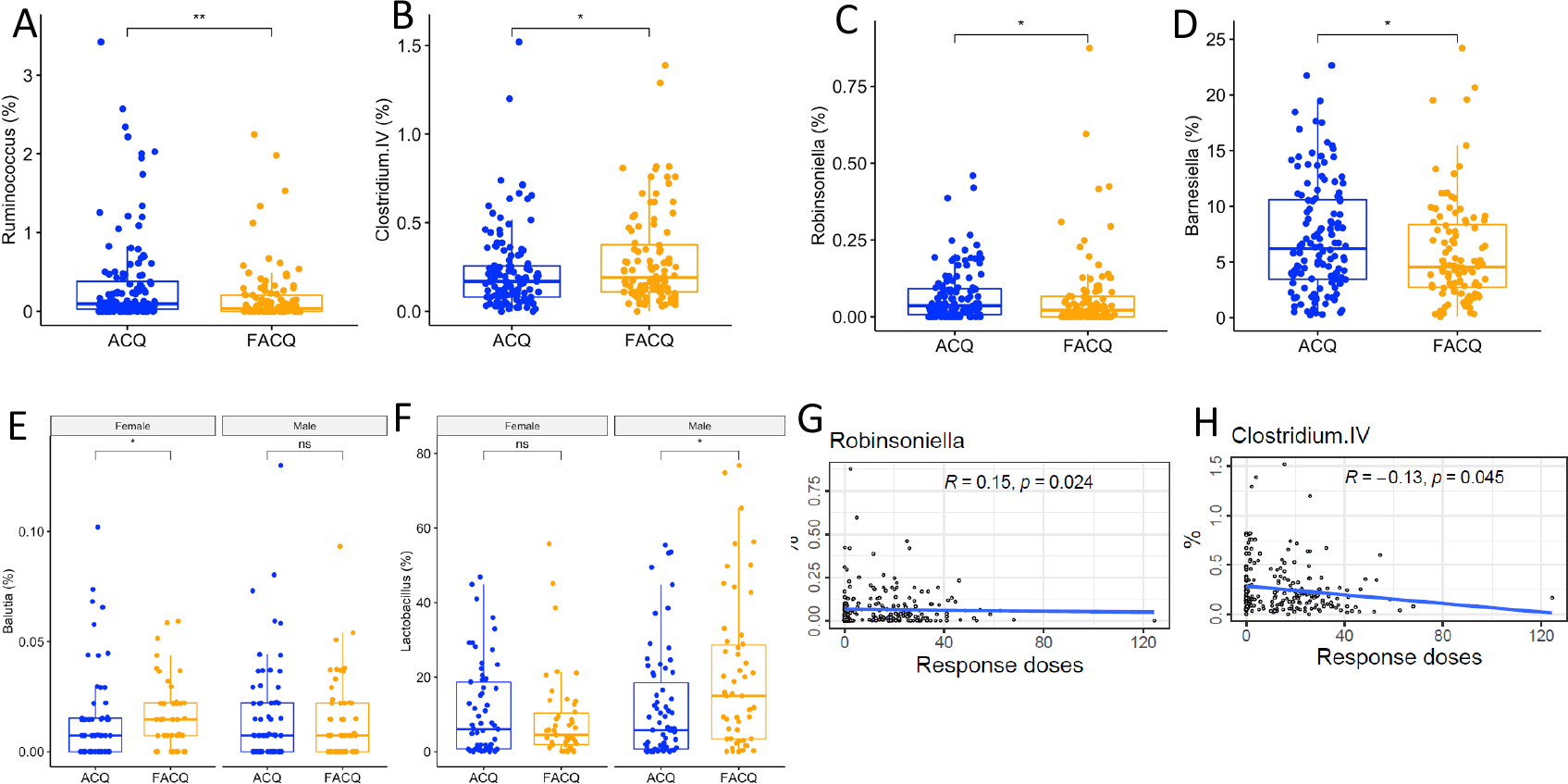
Differential gut microbiome composition of ACQ and FACQ mice. **A)** *Barnesiella*, ***B****) Ruminococcus*, **C)** *Clostridium* IV, **D)** *Robinsoniella*, **E)** Sex-dependent associations *Lactobacillus* and **F)** *Blautia*. **G)** Correlation between *Robinsoniella abundance* and doses of cocaine administered during the acquisition phase. **H)** Correlation between *Clostridium* IV abundance and doses of cocaine administered during the acquisition phase.

**Figure 4.**
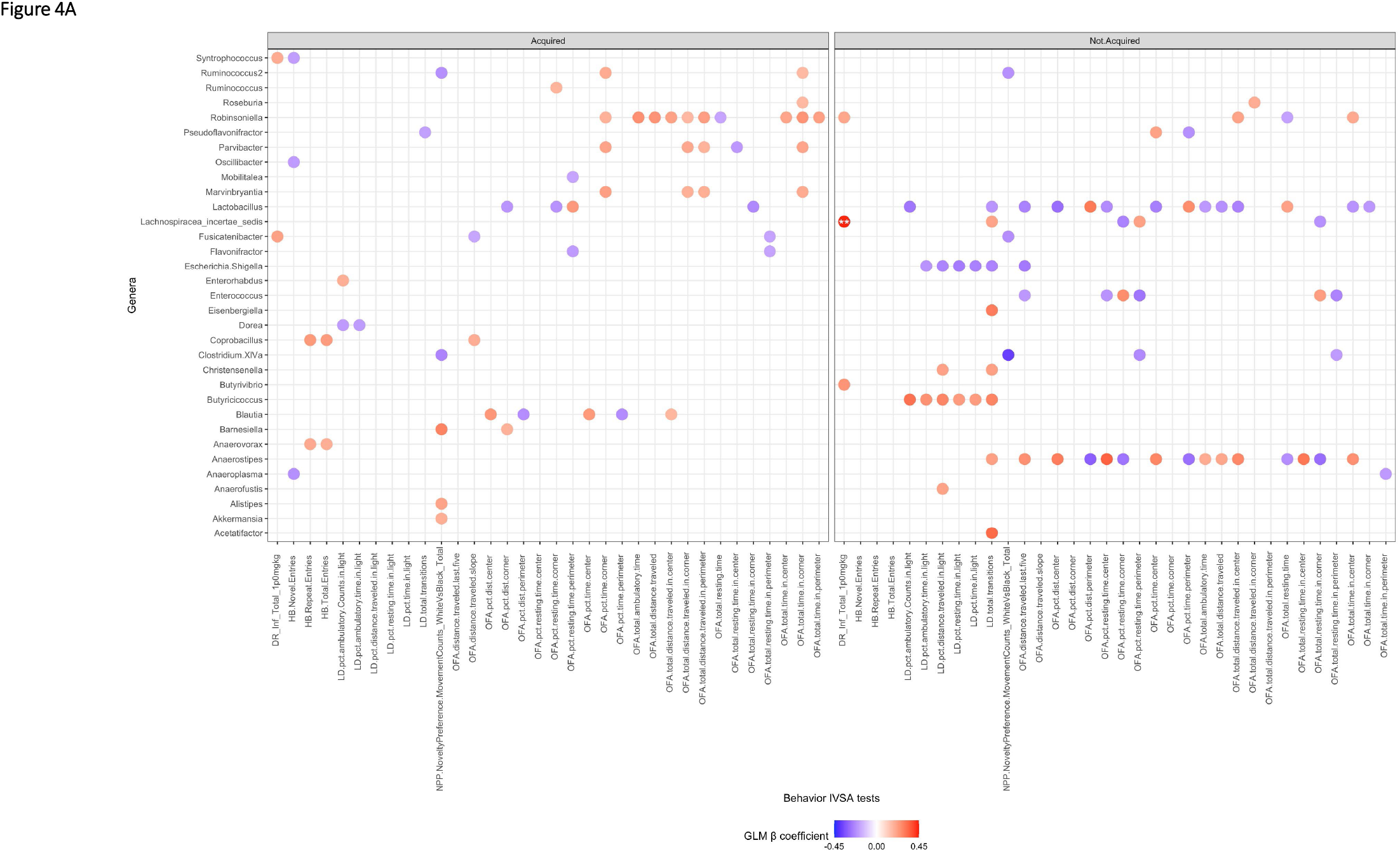
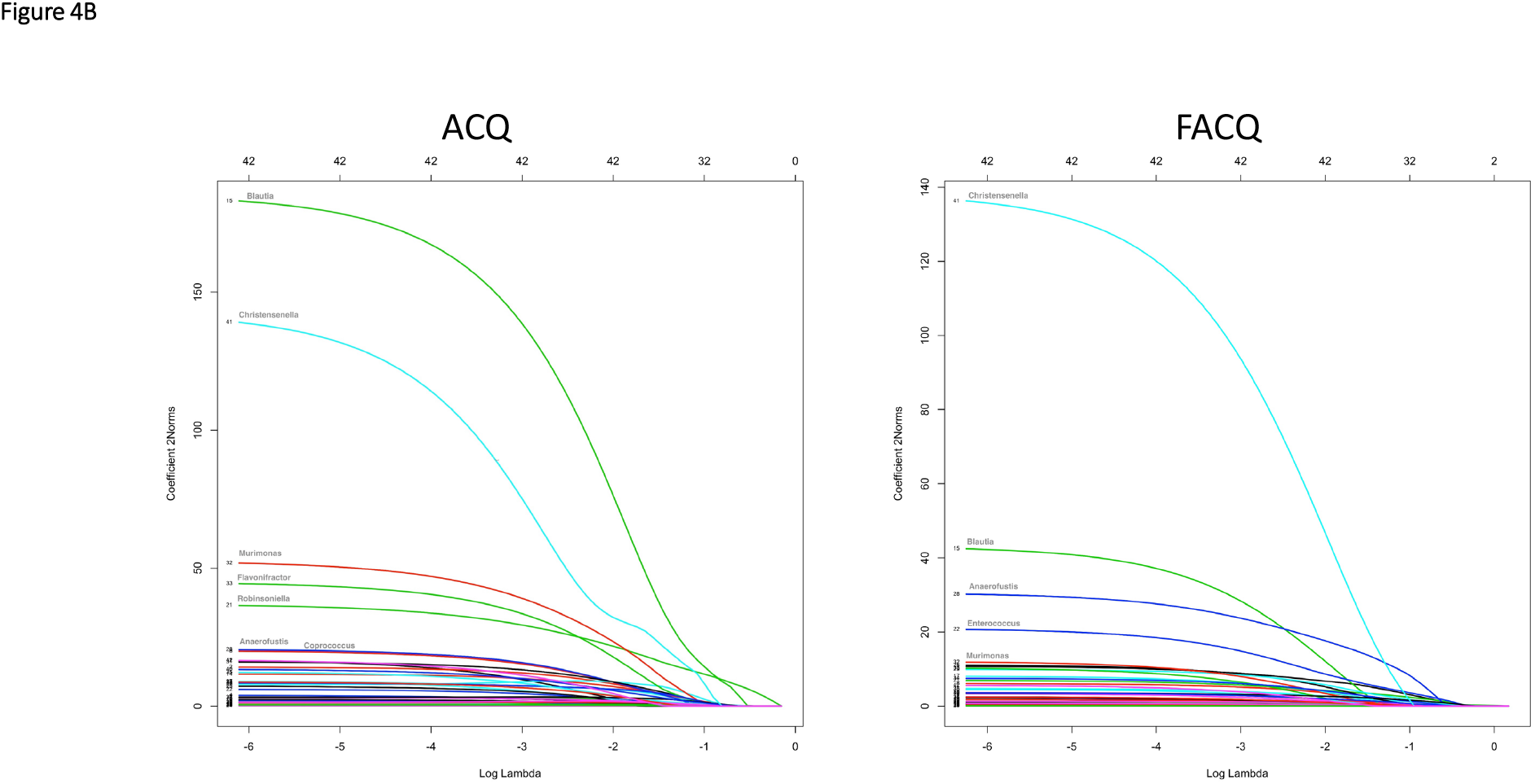
Novelty behaviors associated with microbial abundance in the DO mice. A. Generalized linear model results for specific effects of the microbiome on behavior in DO mice. The GLM fit was Behavior_i_ ~ Age + Sex +Genus_j_ **B)** Lasso regression analysis of the microbiomes as predictors of behavior

### 3.5 Association of the microbiome and behavior in ACQ group

To understand the relationships between the gut microbiome and an array of novelty behaviors of DO mice, we performed GLM with age and sex covariates. Because the microbiome was significantly different between ACQ and FACQ groups, GLM was performed in ACQ and FACQ groups separately. In ACQ group, 57 significant associations were observed (*p*<0.05), but none was significant after p value corrections for multiple testing (**Fig. 4A, Sup Table 3)**. However, *Barnesiella* had the highest regression coefficient with the novelty preference place test (0.28, p=0.001), such that increased *Barnsiella* was associated with increased preference for novel side or novelty. *Robinsoniella* had the largest number of associations with ten open field tests, such that increased activity and decreased anxiety were associated with increased *Robinsoniella* abundance. *Blautia* and *Parvibacter* had five significant associations with behavior in the open field tests, while *Lactobacillus* and *Marvinbryantia* had four significant associations with open field assay. Because no associations passed FDR correction, we sought to validate those findings using LASSO analysis. When carrying out the Lasso regression analysis with all genera, ten genera were found to be associated with thirty-five novelty behaviors and these genera were all *Clostridia* class (*Firmicutes* phylum) (**Fig 4A**). The seven most influential bacteria were *Blautia, Christensenella, Murimonas, Flavonifractor, Robinsoniella, and Anaerofustis*, with the biggest effects on the behaviors in ACQ mice (L2 sum coefficients of 182.9, 138.7, 51.7, 44.3, 36.5, and 20, respectively) following *Coprococcus* (19.8).

### 3.6 Association of microbiome and behavior in the FACQ group across methods

There were 71 significant associations (p<0.05) between the gut microbiome and behaviors in FACQ group, which is larger than the number of associations in the ACQ group (57), although none passed a stringent FDR. (**Fig 4B, Sup Table 3**). *Lactobacillus* and *Anaerostipes* had the largest number of significant associations and opposite associations. An increase in *Lactobacillus* was associated with decreased values in the light-dark and open field, suggesting that FACQ mice are less active and more anxious in the presence of *Lactobacillus. Butyricicoccus, Enterococcus, Escherichia/Shigella, Lachnospiracea_incertae_sedis* and *Robinsoniella generated* significant correlations with the novelty-related behavior tests. Interestingly, different from other microbes, *Butyricicoccus* and *Escherichia/Shigella* were the only two that predominantly correlated to behaviors in the light-dark assay. However, *Butyricicoccus* was positively associated with the light-dark tests, while *Escherichia/Shigella* was negatively correlated to these behaviors. LASSO analysis revealed that the top ten genera that were most important predicted by Lasso regression were all *Clostridia* class (**Fig. 4B**). Interestingly, the coefficient of regression was importantly lower in FACQ than in ACQ mice. For example, *Christensenella* was the most influential microbe, with a coefficient of 116.5 in FACQ mice compared to 138.8 in ACQ mice.

### 3.7 Common and unique associations between the microbiome and behaviors in ACQ and FACQ groups

A common bacterium that repeatedly had significant associations with novelty behaviors in both groups with different statistical methods is the *Robinsoniella* genus. Common associations in both groups reflect similar predicted microbial gene functions, including those involved in metabolic pathways. LASSO analysis also revealed several common short-chain fatty acid-producing bacteria associated with novelty behaviors in both groups. Among those microbes already identified associated with novelty behaviors through GLM additionally *Clostridium XlVb, Clostridium IV*, and *Intestinimonas* were associated through LASSO. In GLM, there are several identical associations between a second Ruminococcus strain (*Ruminococcus2), Clostridium IV* and novelty preference place, *Robinsoniella* and three open field tests (total distance traveled in the center, total resting time, and total time in the center) across groups.

Among 42 analyzed genera, 15 uniquely correlated to novelty behaviors in ACQ mice were mostly from *Clostridia* class such as *Coprococcus, Anaerovorax, Lactonifactor, Ruminococcus, Oscillibacter, Clostridium lV, Syntrophococcus, and Intestinimonas*, these others were from different classes, such as *Barnesiella* and Alistipes (*Bacteroidia* class), *Erysipelotrichaceae_incertae_sedis*, and *Coprobacillus* (*Erysipelotrichia* class), *Akkermansia* (*Verrucomicrobiae* class) *and Parvibacter* (*Actinobacteria* class). Lasso analysis identified *Coprococcus* as an influential microbe of novelty-related behaviors in ACQ mice. Its abundance was negatively correlated to most behaviors in the open field and the light-dark assays. In mice that FACQ, we found only three microbes uniquely correlated to behaviors, two of which belonged to the *Clostridia* class (*Anaerotruncus, Lachnospiracea_incertae_sedis*) and one from the *Gammaproteobacteria* class (*Escherichia*.*Shigella*).

### 3.8 Network analysis shows complex interactions between bacteria abundance and novelty behaviors and the response to the number of infusions of the cocaine self-administered, revealed important association between Coprococcus and Actinobacteria phylum in FACQ

The network prototype was built on filtered microbial data (filter of 50% of OTU prevalence), and all novelty behaviors were integrated, with each novelty behavior being considered a node like an OTU (**Fig 5A**). The microbes that are consistently present seemed to be similar between ACQ mice and FACQ mice. The network presented 241 nodes and 342 edges for ACQ mice, while the FACQ mice network had 243 nodes and 346 edges. The interactions in both networks were more likely dominated between OTUs of the highest dominant phylum *Firmicutes* (195 connections in ACQ mice, 199 connections in FACQ mice) and between OTUs of Bacteroidetes (43 connections in ACQ mice, 47 connections in FACQ mice). Moreover, the interactions between OTUs of *Firmicutes*-*Bacteroidetes* and between OTUs of *Firmicutes*-*Actinobacteria* were higher in FACQ mice (21 and 21 respectively) than in ACQ mice (17 and 13 respectively). Among the interactions between microbial taxa and novelty behaviors, it is interesting to note that *Coprococcus* (OTU_116) (also identified in the LASSO analysis) and *Lactobacillus* (OTU_1) were the two common bacteria correlated to several novelty tests with an almost inverse relationship. However, in ACQ mice, *Coprococcus* OTU was negatively associated with percent resting time in the light (LD3), percent resting time in the perimeter (OFA21) and *Clostridium IV* (OTU_193). In FACQ mice, *Coprococcus* (OTU_116) was negatively correlated to novelty place preference (NPP1), percent distance in the center (OFA13) and percent resting time in the corner (OFA20) of open field assay. We also observed the positive associations between *Lactobacillus* (OTU_1) and behaviors in open field assay in both groups, such as positively with percent distance in the perimeter (OFA15) for FACQ mice, positively with percent resting time in the perimeter (OFA21) for ACQ mice. Infusions at 1.0mg/kg (IVSA) was associated with percent resting time in the center (OFA19) in FACQ and with percent time in the corner (OFA17) and percent resting time in the corner (OFA20) in ACQ mice. The important main difference between the two networks remains the negative association between *Coprococcus* (OTU_116) and *Enterohardus* (OTU_58, *Actinobacteria* phylum), an important node in the network that generated ten connections with mostly all other species of *Actinobacteria* phylum (*Enterohardus* (OTU_7, OTU_76), *Parvibacter* (OTU_229), *unclassified_Coriobacteriaceae* (OTU_24, OTU_249, OTU_208, OTU_33). This association was absent in ACQ mice.

**Figure 5.**
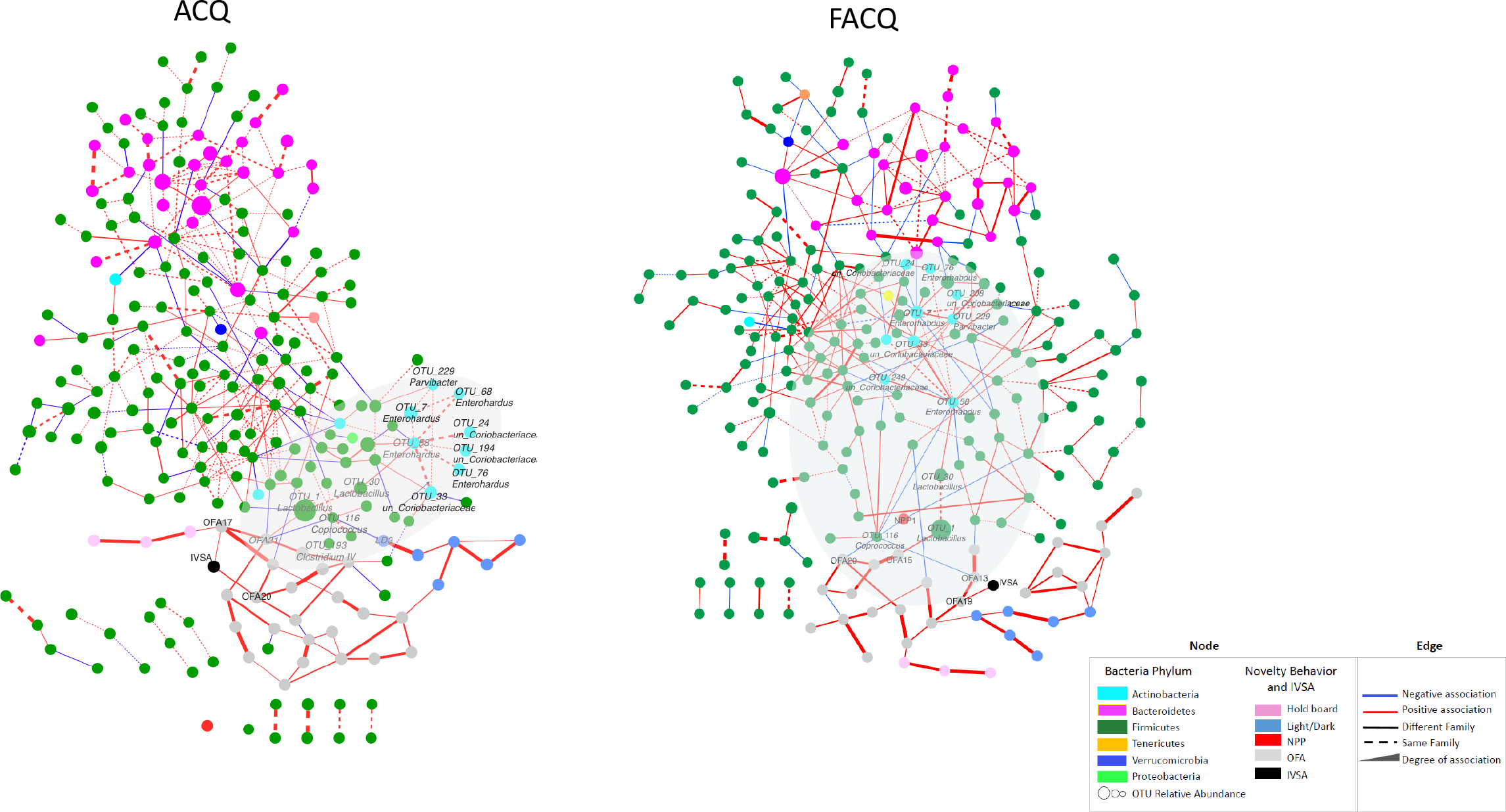
Network analysis of ACQ and FACQ at OTU level: interactions between microbe-microbe and microbe-behavior. The network was built on filtered 50% OTU prevalence of DO mice cohort using SPIEC-EASI. Each OTU and each novelty behavior was considered as a node in the network. Abbreviations in network: novelty place preference (NPP1), percent distance in the center (OFA13), percent distance in the perimeter (OFA15), percent time in the corner (OFA17), percent resting time in the center (OFA19), percent resting time in the corner (OFA20), percent resting time in the perimeter (OFA21), percent resting time in the light (LD3), Number of infusions at 1.0 mg/kg (IVSA).

### 3.9 Functional profiling and mWGS analysis show various aspects of glutamate metabolism

Using PICRUSt2 on the 16S data, we identified 32 gut-brain modules as significantly (p<0.05) different between the microbiome of ACQ and FACQ mice and two with qvalue <0.05 (**Fig 6 A-D, Table 1**). Five modules mapped to the GABA I synthesis KO, 18 to glutamate, and seven to propionate degradation/synthesis, in addition to one to dopamine and gamma-hydroxybutyric acid degradation KO (GHB). From mWGS data on a subset of 96 mice, we identified eight gut-brain modules that differed significantly (p<0.05) between ACQ and FACQ group (**Table 2**). Half of those modules were related to glutamate, while the others were methionine biosynthesis, acetate synthesis I, isovaleric acid synthesis I and S-adenosylmethionine synthesis. Both methodologies identified other-glutamate modules KO1919 and KO10004 as significantly up-regulated in mWGS and 16S. These glutamate KO were glutamine-cysteine ligase and glutamate/aspartate transport system ATP binding protein respectively. Pathway analysis of the mWGS data showed three pathways upregulated greater than 0.5 fold, (glutamate degradation PWY-4321, peptidoglycan biosynthesis IV PWY-6471, heme biosynthesis II HEMESYN2.PWY) and one pathway, fatty acid salvage PWY-7094, which were downregulated at least 0.5 fold in FACQ mice. (**Fig 6F, Table 3**). The source of the PWY-4321 gene produced from the mWGS sequencing data indicated that the *Lactobacillius reute*ri was one of the sources of these pathway members (**Sup Table 4**). We looked at the three most abundant species of *Lactobacillus (L*.*reuteri, L. gasseri, L. intestinalis*) and found that they were significantly (p<0.05) increased in FACQ mice compared to ACQ mice (**Fig 6G**). These species are well known for their immunomodulatory effects on the central nervous system and anxiety-like behavior. Furthermore, *Lactobacillus* species are known to produce GABA and glutamate^8^. Together functional profiling of 16S and mWGS data showed enriched trends toward glutamate biosynthesis in the FACQ group with the results showing remarkable agreement between the two sequencing types.

**Table 1.**
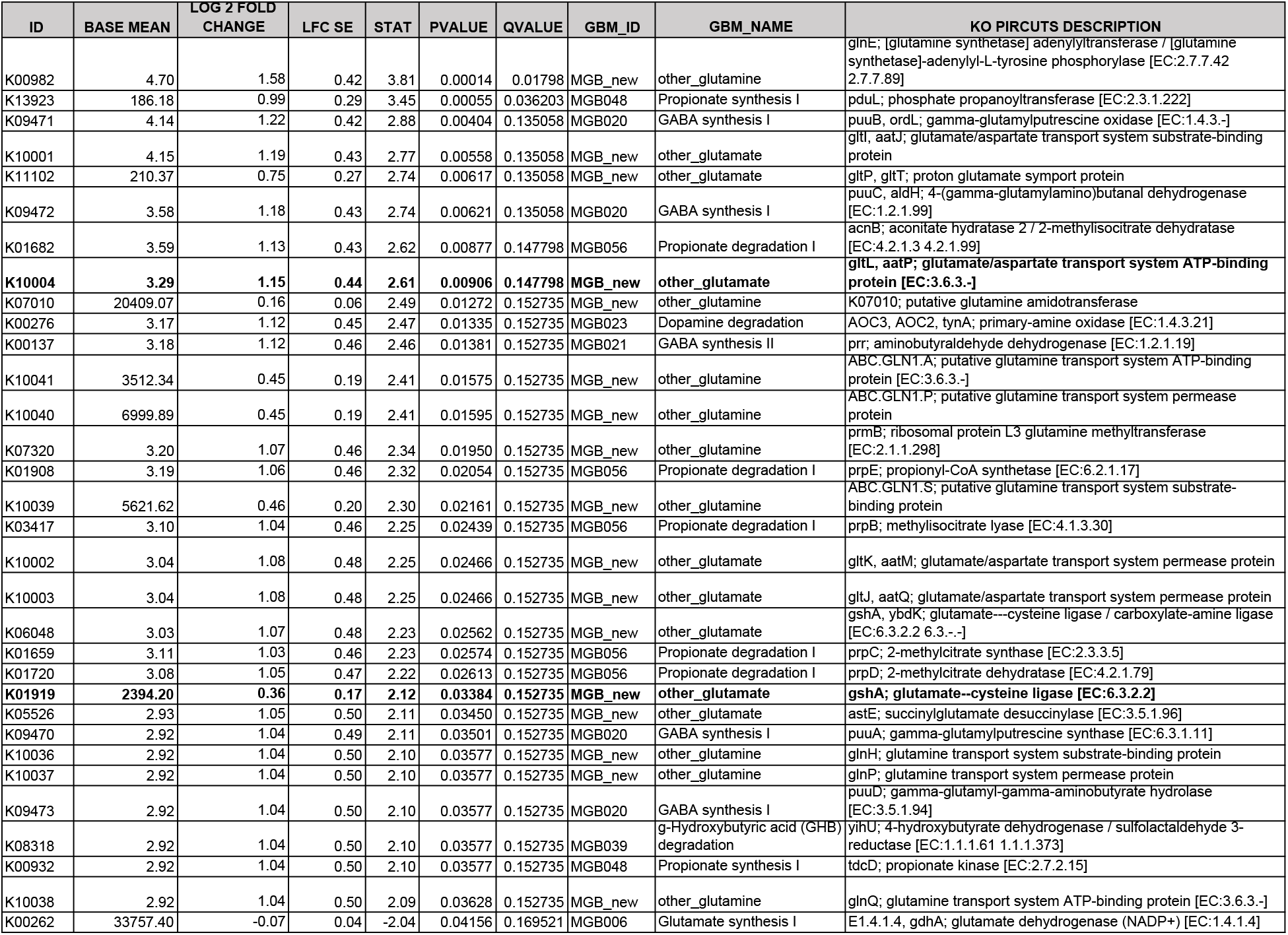
Differential Metabolic Potential from PICRUSt of 16S data.

**Table 2.** Differential Metabolic Potential from mWGS.

| ID | BASE MEAN | LOG 2 FOLD CHANGE | LFC SE | STAT | PVALUE | QVALUE | GBM_ID | GBM_NAME | KO.name |
| --- | --- | --- | --- | --- | --- | --- | --- | --- | --- |
| K01928 | 277.38 | 0.14 | 0.05 | 3.01 | 0.00259 | 0.25929 | MGB_new | other_glutamate | K01928: UDP-N-acetylmuramoylalanyl-D-glutamate--2,6-diaminopimelate ligase [EC:6.3.2.13] |
| K00549 | 6.58 | 1.44 | 0.50 | 2.90 | 0.00379 | 0.25929 | MGB_new | Methionine biosynthesis, apartate => homoserine => methionine | K00549: 5-methyltetrahydropteroyltriglutamate--homocysteine methyltransferase [EC:2.1.1.14] |
| K01951 | 321.30 | 0.16 | 0.06 | 2.52 | 0.01169 | 0.51059 | MGB_new | other_glutamine | K01951: GMP synthase (glutamine-hydrolysing) [EC:6.3.5.2] |
| **K01919** | **14.12** | **0.78** | **0.32** | **2.43** | **0.01491** | **0.51059** | **MGB_new** | **other_glutamate** | **K01919: glutamate--cysteine ligase [EC:6.3.2.2]** |
| **K10004** | **46.86** | **0.47** | **0.20** | **2.33** | **0.01999** | **0.52988** | **MGB_new** | **other_glutamate** | **K10004: glutamate/aspartate transport system ATP-binding protein [EC:3.6.3.-]** |
| K00625 | 252.35 | 0.11 | 0.05 | 2.23 | 0.02543 | 0.52988 | MGB043 | Acetate synthesis I | K00625: phosphate acetyltransferase [EC:2.3.1.8] |
| K01073 | 2.35 | 1.30 | 0.59 | 2.21 | 0.02707 | 0.52988 | MGB034 | Isovaleric acid synthesis I (KADH pathway) | K01073: acyl-CoA hydrolase [EC:3.1.2.20] |
| K00789 | 488.20 | 0.11 | 0.05 | 2.12 | 0.03433 | 0.58795 | MGB036 | S-Adenosylmethionine (SAM) synthesis | K00789: S-adenosylmethionine synthetase [EC:2.5.1.6] |

**Table 3.**
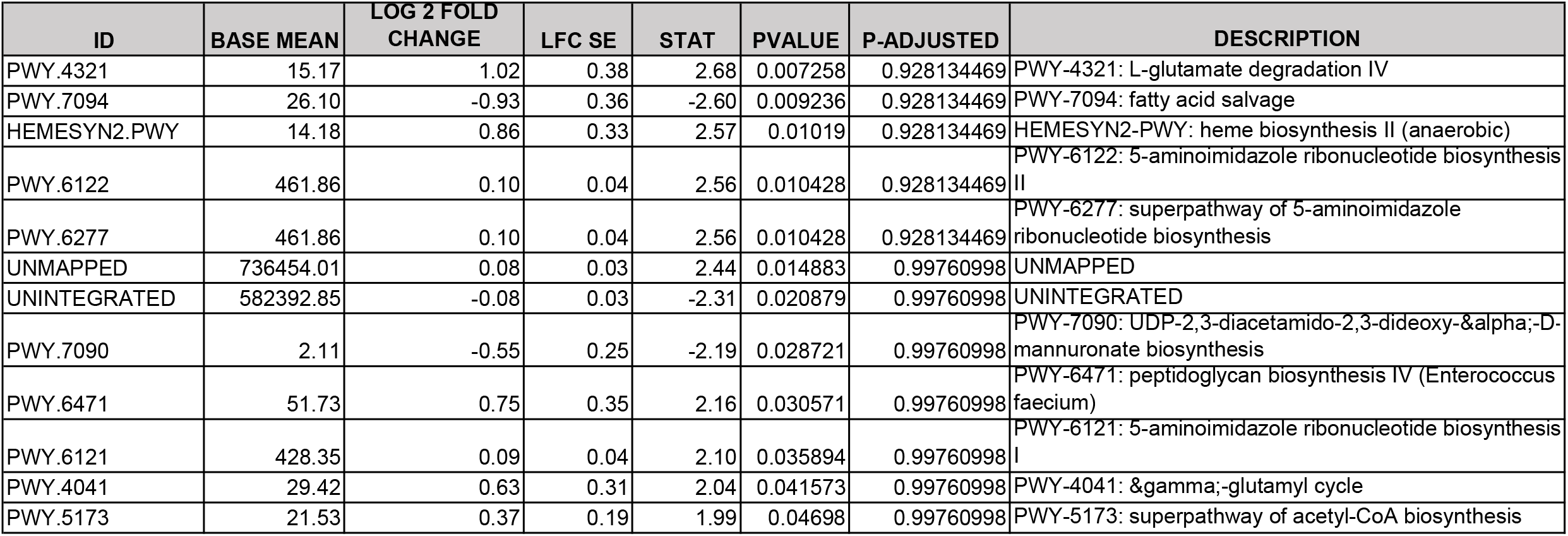
Differential Pathway Abundance from WGS.

**Figure 6.**
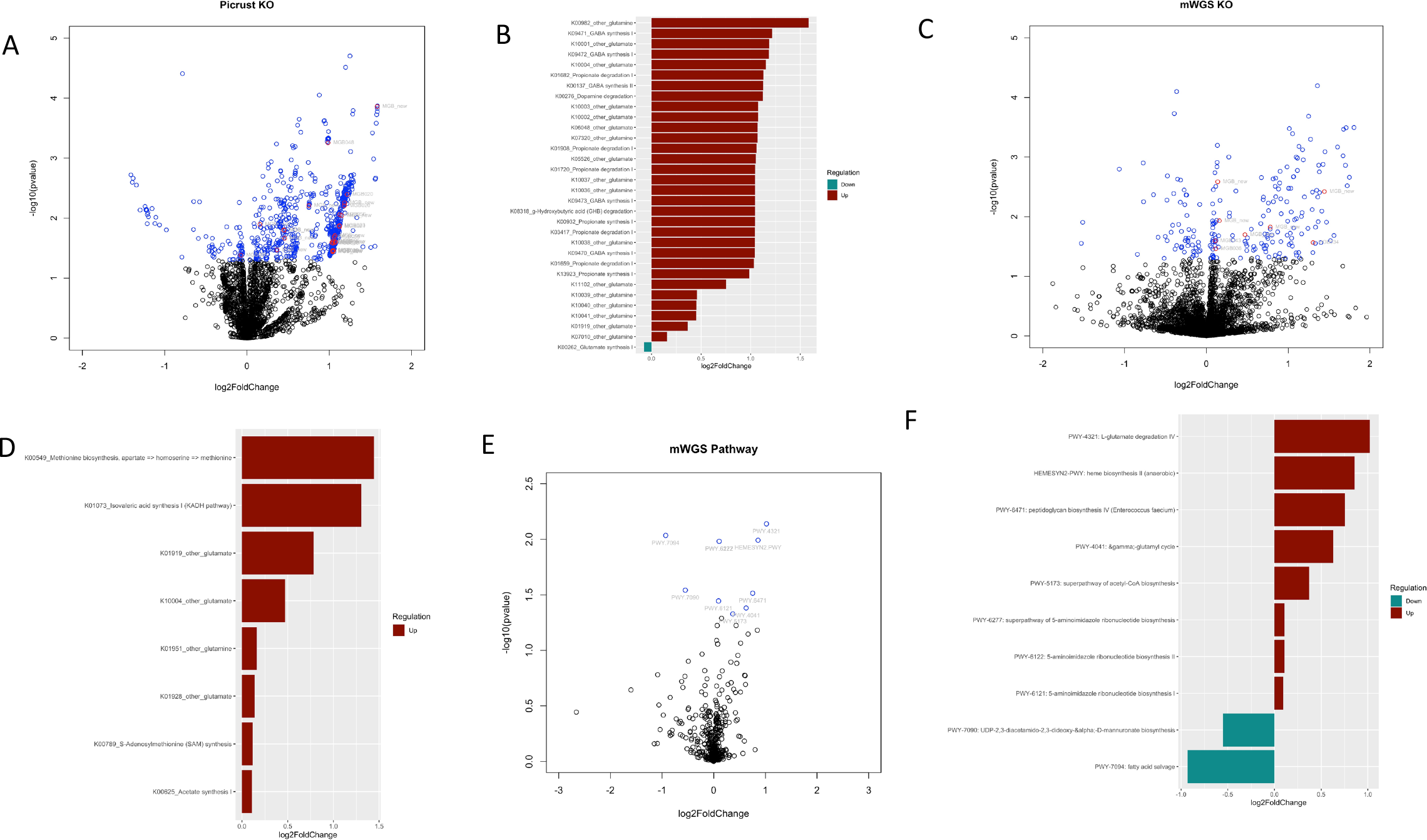
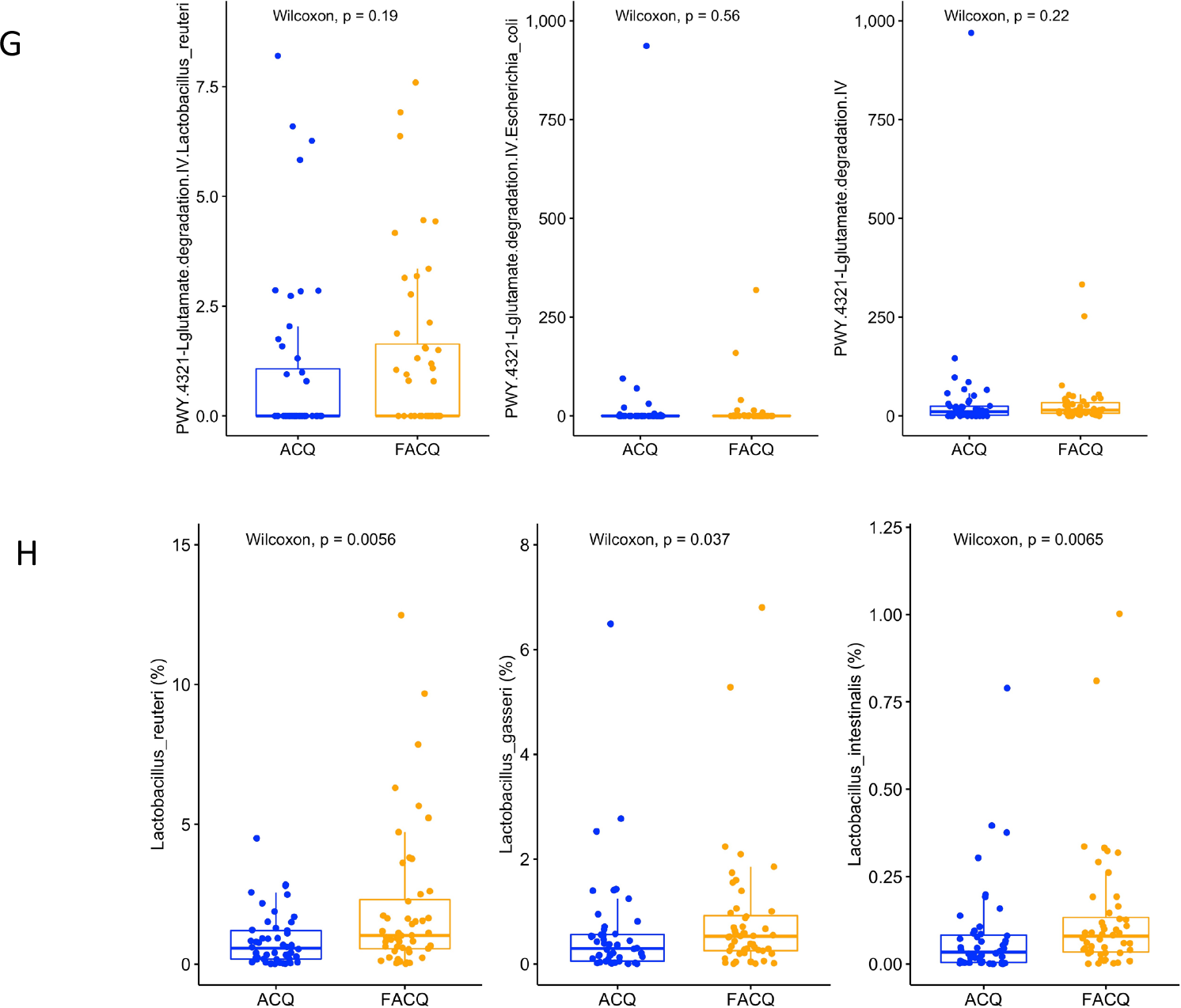
Volcano plots of the functional categories encoded by the microbiome. **A**. A scatterplot showing the statistical significance and magnitude of change of the KO clusters as determined by PICRUSt2 of 16S data. **B**. The KO categories (p<0.05) from PICRUSt2 of 16S data. **C**. A scatterplot of the statistical significance and magnitude of change of the KO clusters determined by WGS. **D**. The KO categories (p<0.05) from WGS data. **E**. A scatterplot of the statistical significance and magnitude of change of the mWGS Pathways **F**. The mWGS Pathways (p<0.05) Blue=significant KO/pathway (p value<0.05), red=significant KOs that are GBM (p value<0.05). **G. H**.

## DISCUSSION

Growing evidence suggests that understanding the complex relationship between novelty-related behaviors, intravenous self-administration and gut microbiome composition is crucial in shedding light on the factors associated with gut-brain axis communication and addiction and addiction-like behaviors. Our study presents a large microbiome dataset from the feces, novelty-related behavior assays and intravenous self-administration to assess the bi-directional gut-brain axis communication in a random population of DO mice. The associations provide a better perspective of gut-brain axis communication and reflect predicted microbial gene function, including those involved in glutamate pathways. Our profiling of the microbiome and IVSA identified an association of four genera of bacteria *Barnesiella, Ruminococcus, and Robinsoniella* abundance positively correlated with acquisition and *Clostridium* IV negatively correlated with the acquisition of self-administration behavior. In addition, we found a sex-specific association of *Lactobacillus* abundance with the ACQ phenotype in males and *Blautia* abundance in females with the ACQ phenotype. Further analysis of the metabolic pathways encoded by the microbiome as measured by both 16S and mWGS supports a key differential role for the microbiome in glutamate metabolism and evidence for its role in dopamine degradation. These results are interesting in the context of the neurotransmitters in the brain, which we know are associated with addiction^50–53^.

In previous studies, the abundance of *Barnesiella*, an SCFA-producing microbe, was increased tenfold in colonic contents in response to cocaine.^54^ In the same study, *Pseudoflavonifractor* was decreased by cocaine administration. Here we see *Barnesiella* correlated with novelty preference and percent center time in the ACQ group by the GLM approach. *Pseudoflavonifractor* was associated with the number of light-dark transitions in the ACQ group and two open field measures in the FACQ group. In outbred rats, the cecal *Barnesiella* abundance was negatively correlated with locomotor activity in a sex-specific fashion.^55^ *Fusicatenibacter* here in both groups was associated with novelty preference and, in a human study, was one of the taxa related to individuals undergoing a depressive episode^56^

The role of the microbiome in behavior has been well established. Germ-free (GF) mice exhibit abnormal immune- and neuro-development and display abnormal behaviors such as increased locomotor activity in the open field^57^. The behavioral abnormalities, reflecting decreased anxiety, extend to time spent in the center of the open fields and increased time spent on the light side of the light-dark apparatus^58^. The changes observed in the GF mice, specifically the open field center time and center distance, were shown to not be reversible by colonization with microbiota from a specific pathogen-free facility^59^. However, probiotic treatment of GF mice with live *Lactobacillus plantarum* PS128 resulted in increased total distance traveled and decreased center time in the open field. Recent findings have even shown that a gut-derived metabolite such as 4-ethylphenyl sulfate influences anxiety-like behaviors in mice through alteration of oligodendrocyte function and myelin pattering in the brain.^60^ Transplant studies into mice from people with schizophrenia vs. controls resulted in increased distance traveled in the open field and increased center time. This behavior change suggested that the dysbiosis seen in schizophrenia can function through modulating the glutamate-glutamine-GABA cycle in the hippocampus^61^. The association of *Lactobacillus* and glutamate metabolism with cocaine IVSA acquisition is very interesting. Imaging studies have shown that BALBc/J mice given *Lactobacillus rhamnosus*, although not one of the species identified here, resulted in increases in brain GABA, N-acetyl aspartate and glutamate^62^.

Data transformation greatly influences downstream analyses. Rarefaction^68^ and normalization are well-known methods for working with microbiome data. Observed differences may be affected by variations in sequencing quality rather than true biological differences without rarefaction. However, different normalization approaches may lead to other association structures and clustering results, as well as depend on the nature of the data. Our results relayed largely on logarithms and a high proportion of zeros in the dataset. As a result, prediction and feature selection methods considerably depend on handling zeros.

Here we associate microbial abundance of four genera with the propensity of mice to acquire self-administration of cocaine. Others have shown that within one homogeneous inbred strain, a subset of mice often fails to acquire self-adminsitration^63^. Roberts *et al*. looked at eight different inbred strains and found an acquisition rate anywhere from 100% (I/LnJ) to 47% (FVN/NJ). It is possible that due to cage effects or parent of origin effects, or another environmental variable, these mice may contain a different microbiome than those that do acquire IVSA. We know that diet can have a profound impact on behavior, as was recently illustrated by the Linsenbardt group, which showed that the development of alcohol front-loading was abolished by switching mice C57BL/6J mice from a LabDiet5001 to a Teklad 2920x rodent diet^64^. The role of the microbiome was not addressed in that work, but we know that diet and environment are the biggest drivers of microbiome composition in a genetically inbred population.^65^ Work using the same assays of binge drinking in C57BL/6J mice with a 2-week antibiotic pretreatment significantly increased alcohol consumption^66^, and this increased consumption could be reversed by sodium butyrate supplementation^67^.

This study demonstrates, by using multiple approaches in an outbred mouse population with genetic variation similar to the human population, that certain bacteria are specifically associated with novelty response and its predisposing effects on intravenous self-administration of cocaine. The strongest associations were the relative abundance of *Barnesiella, Ruminococcus, Robinsoniella* and *Clostridium* IV with the ability to self-administer cocaine. In addition, the sex-specific association of *Lactobacillus* with IVSA and the gut-brain modules it is associated with supports a role for microbiomes encoding glutamate metabolism in the ability to self-administer cocaine. This finding provides evidence of alternate pathways that can be modulated to alter addiction behavior through manipulating the microbiome using metabolites or pre-and probiotics.

## ABBREVIATIONS

DO: diversity outbred;
IVSA: intravenous self-administration;
OTU: operational taxonomic unit.
PICRUSt: Phylogenetic Investigation of Communities by Reconstruction of Unobserved States
SCFA: short chain fatty acids;
WGS: whole genome sequencing;
ACQ-Acquired: cocaine self-administration phenotype;
FACQ: (Failed to acquire cocaine self-administration behavior);
KO-KEGG: Orthology, GBM gut-brain modules

## CRediT authorship contribution statement

**Thi Dong Binh Tran:** Methodology, Formal analysis, Data Curation, Investigation, Writing-Original Draft, Writing-Review and Editing, Visualization

**Hoan Nguyen**: Formal analysis, Data Curation, Investigation

**Erica Sodergren**: Project administration, Investigation

**Center for Systems Neurogenetics of Addiction**: Conceptualization, Methodology, Investigation, Formal analysis, Data Curation

**Price E Dickson:** Methodology, Investigation, Supervision, Writing-Review and Editing

**Susan N Wright**: **Writing**-Review and Editing

**Vivek M Philip**: Methodology, Conceptualization, Investigation, Formal analysis, Data Curation, Writing-Review and Editing,

**George M. Weinstock**: Methodology, Conceptualization, Writing-Original Draft, Writing-Review and Editing, Project administration, Funding Acquisition,

**Elissa J Chesler:** Conceptualization, Writing-Review and Editing, Project administration, Funding Acquisition

**Yanjiao Zhou:** Conceptualization, Writing-Original Draft, Writing-Review and Editing, Funding Acquisition,

**Jason A. Bubier:** Conceptualization, Writing-Original Draft, Writing-Review and Editing, Project administration, Funding Acquisition,

## Declaration of competing interest

### Code Availability

The R scripts used for all data analysis available at <GitHub>.

### Data availability

16S Data are deposited at the SRA, accession **#######**. The mouse phenotype data is available in the mouse phenome database in project CSNA03.

## Acknowledgments

NIH R01 DA037927 to EJC; NIH P50 DA039841 to EJC; NIH U01 DA043809 to JAB/GW. Cocaine hydrochloride was provided by NIDA Drug Supply Program.

## Figure Legends

**Table 1-16S metabolic potential analysis**.

**Table 2-WGS metabolic potential analysis**.

**Table 3-WGS Pathway Analysis**

**Supplemental Figure 1-** Microbiome composition of top 25 genera in DO mice.

**Supplemental Figure 2-** Correlation of each microbe with the number of doses of 1.0mg/kg taken during acquisition.

**Sup Table 1**-**The variable names and all the metadata generated by the equipment for Open Field, Light Dark and Holeboard**.

**Sup Table 2**-**Summary table of Wilcoxon rank sum test for the relationship between microbes, behavior, sex and IVSA status (ACQ/FACQ). Pink = p<0.05**

**Sup Table 3-GLM and LASSO data for ACQ and FACQ groups**

**Sup. Table 4-Differential Pathways between ACQ and FACQ by mWGS sequencing**

