## Supplementary figures and images for "Microbial glutamate metabolism predicts intravenous cocaine self-administration in Diversity Outbred mice"

### Sup Fig 1

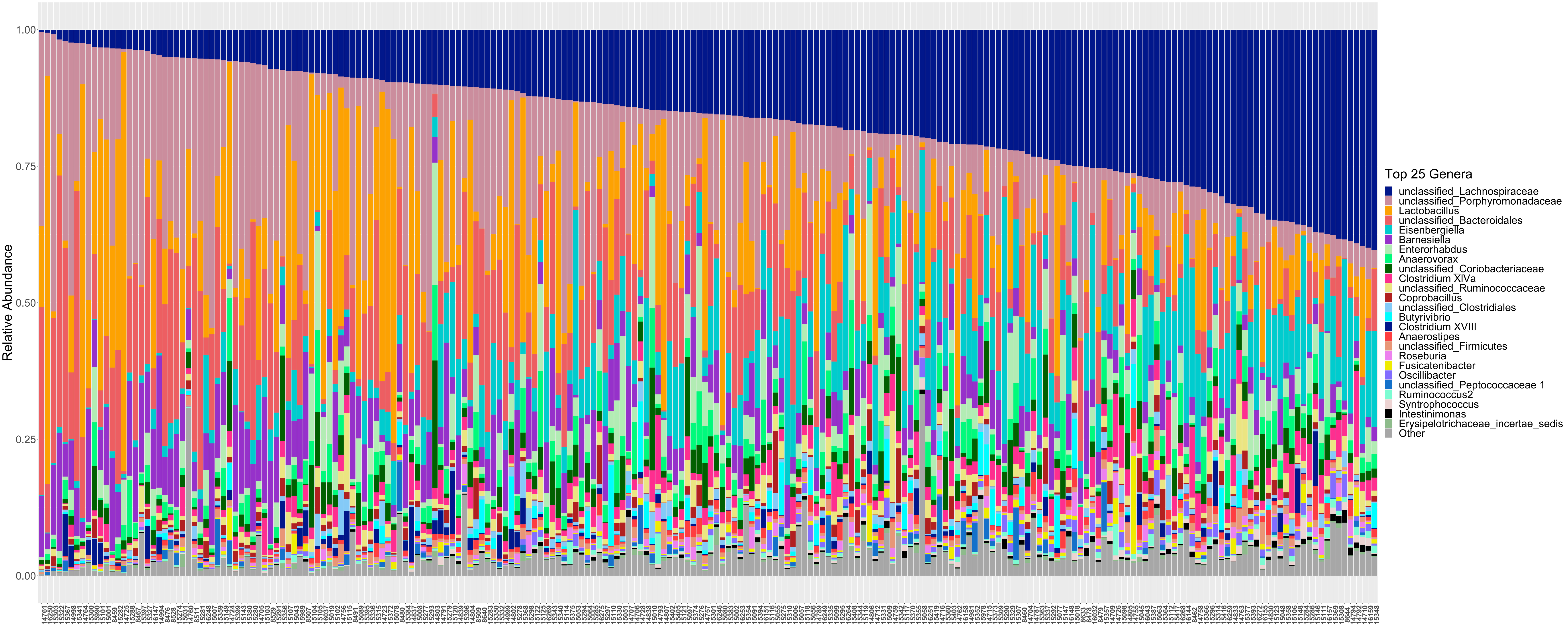

### Sup Fig 2

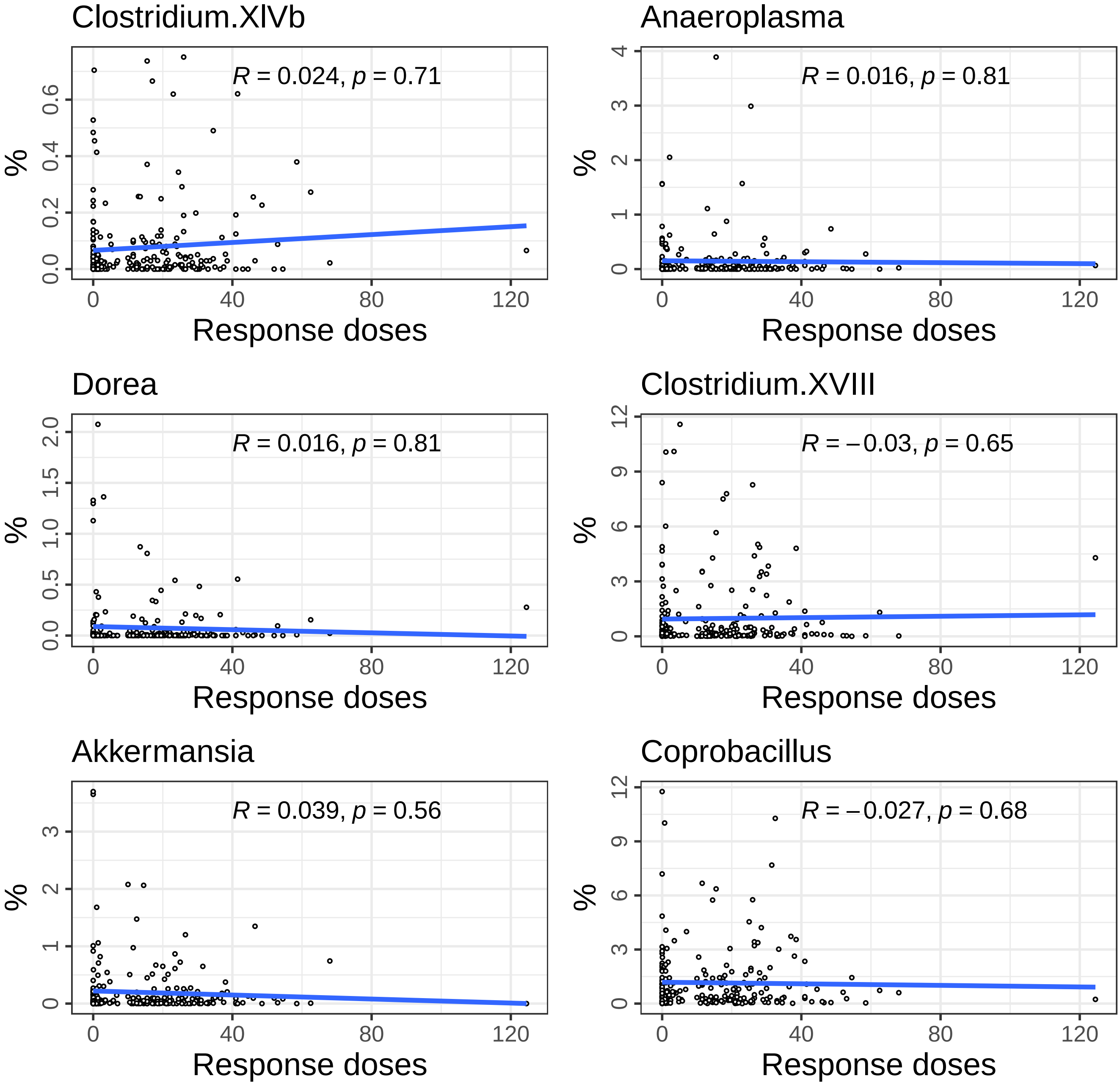

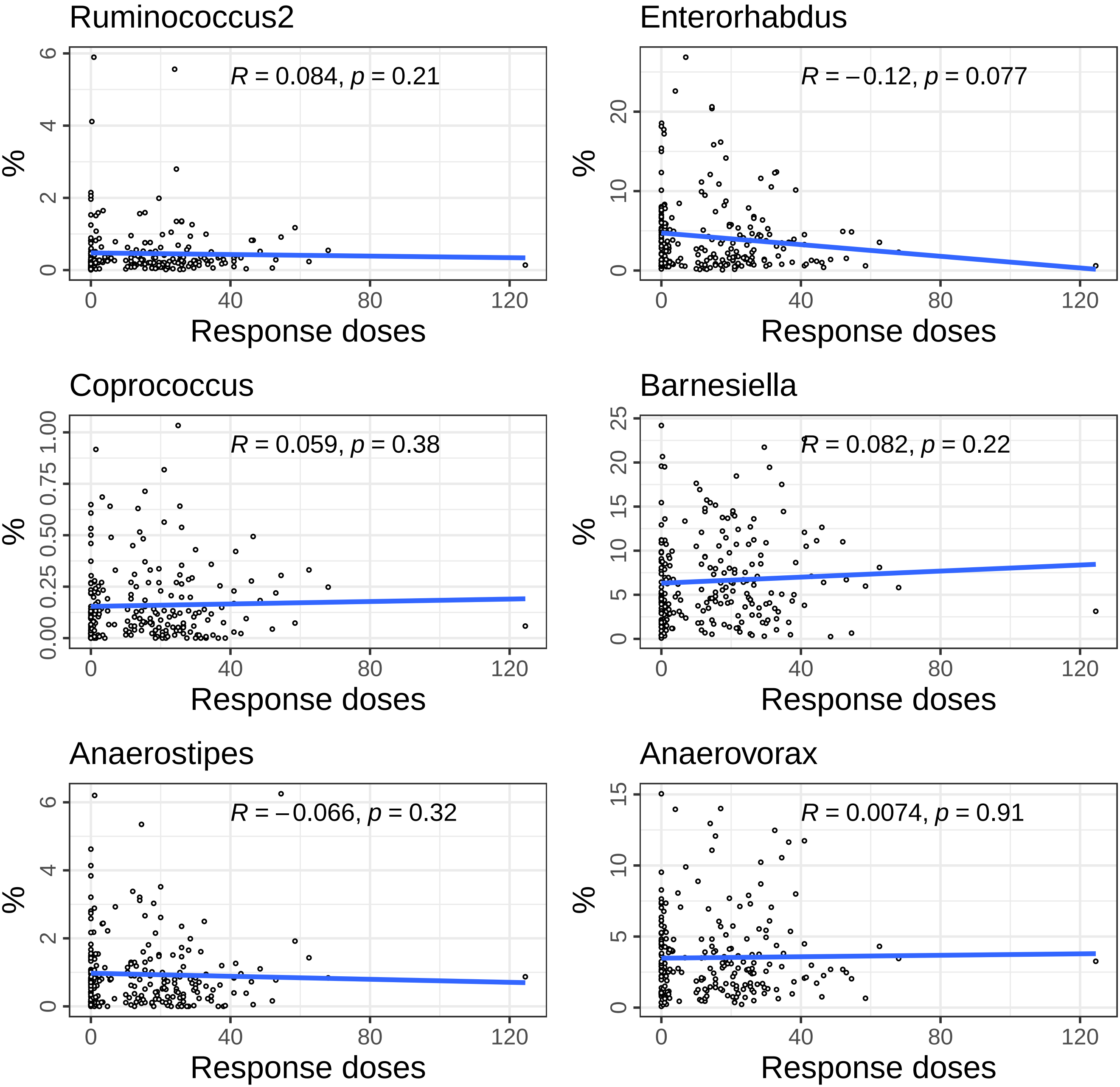

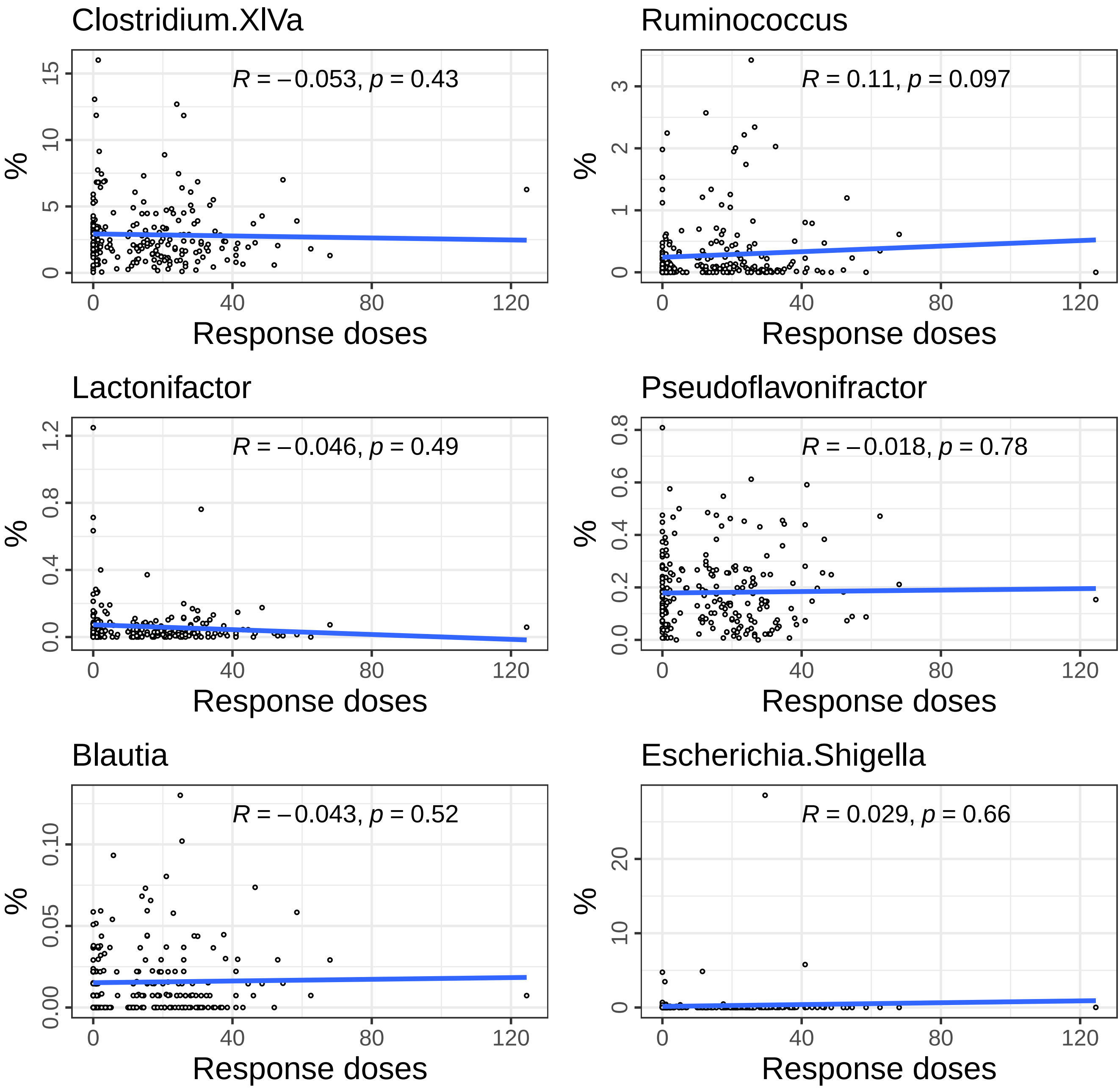

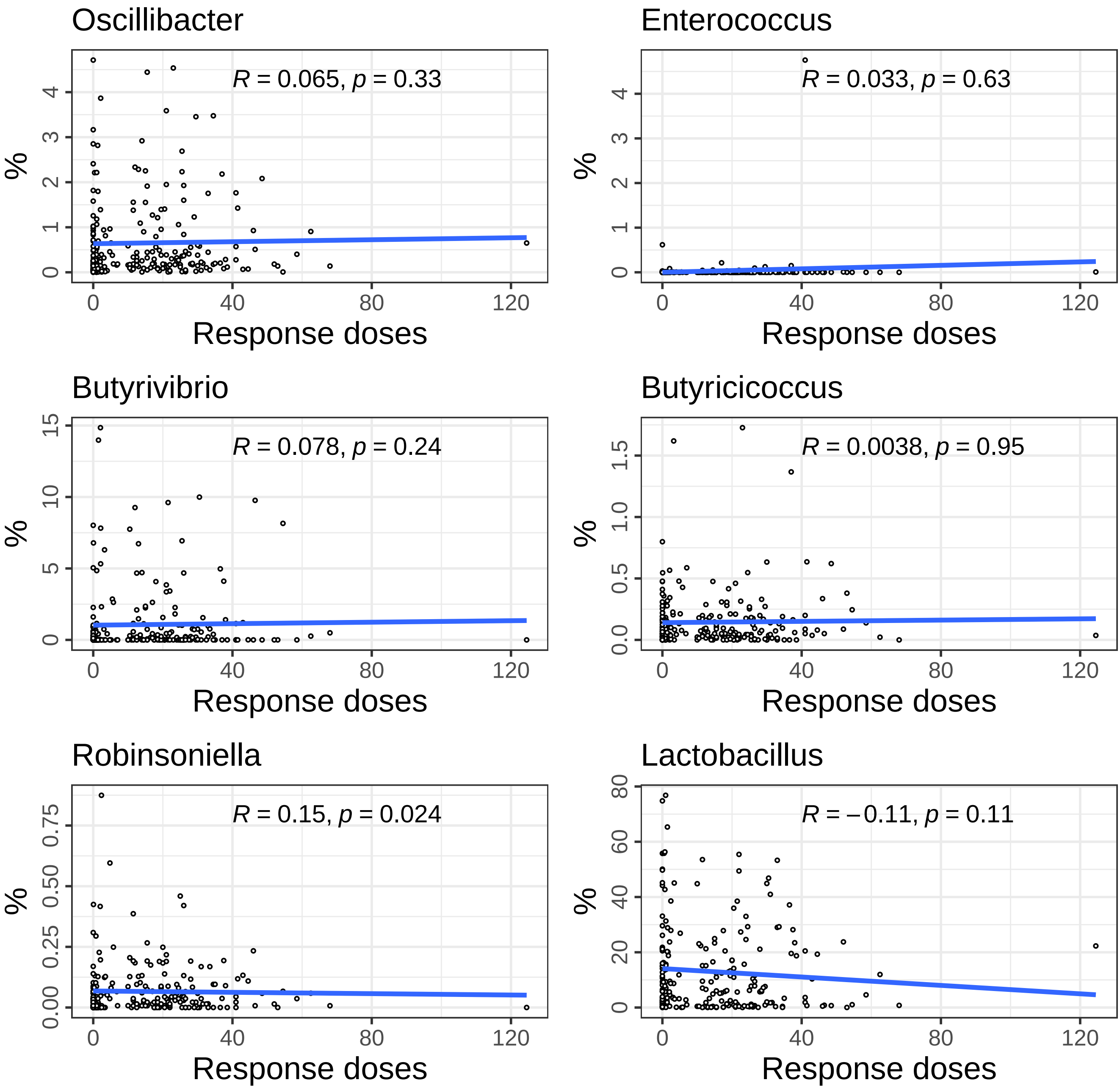

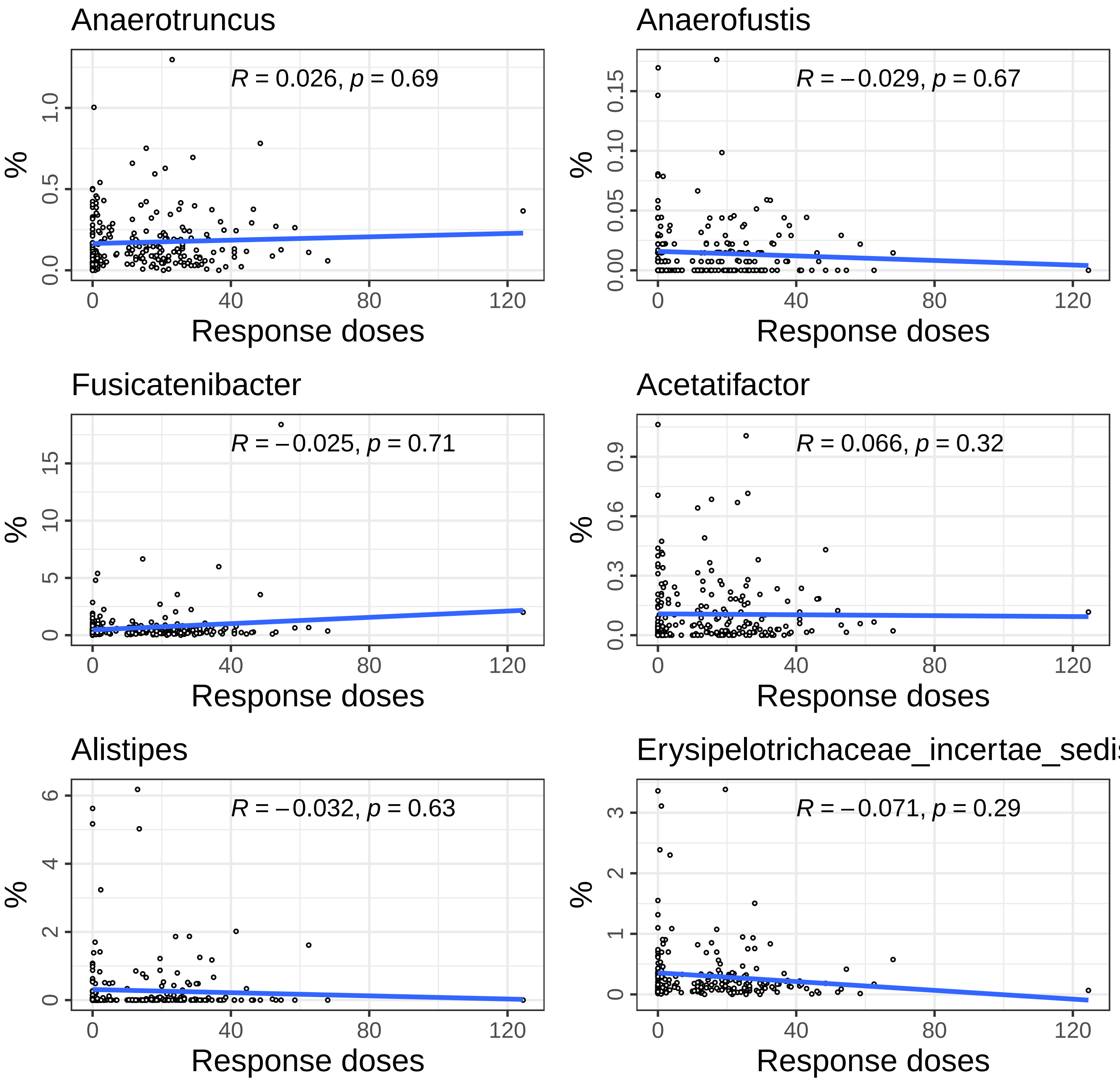

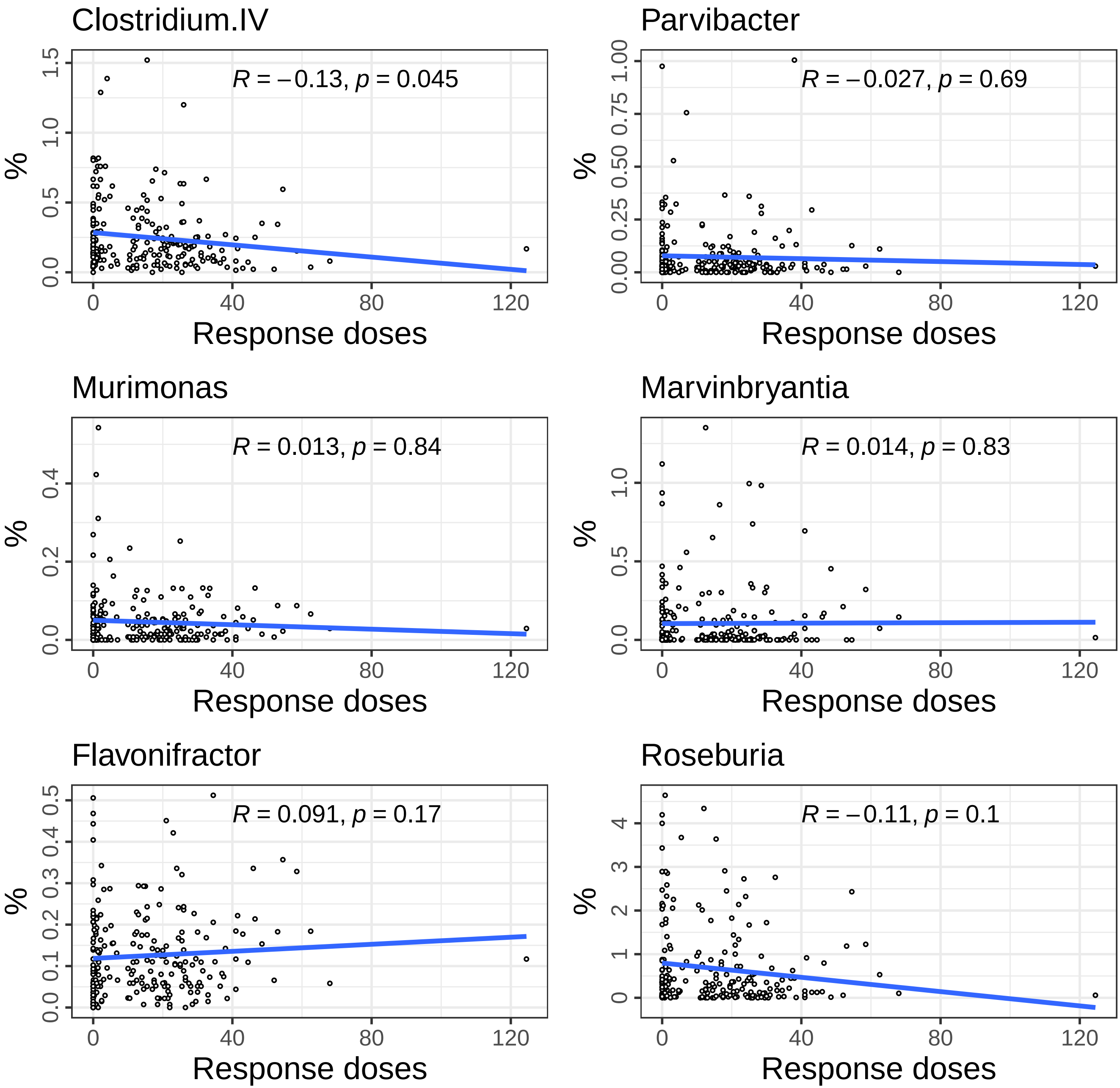

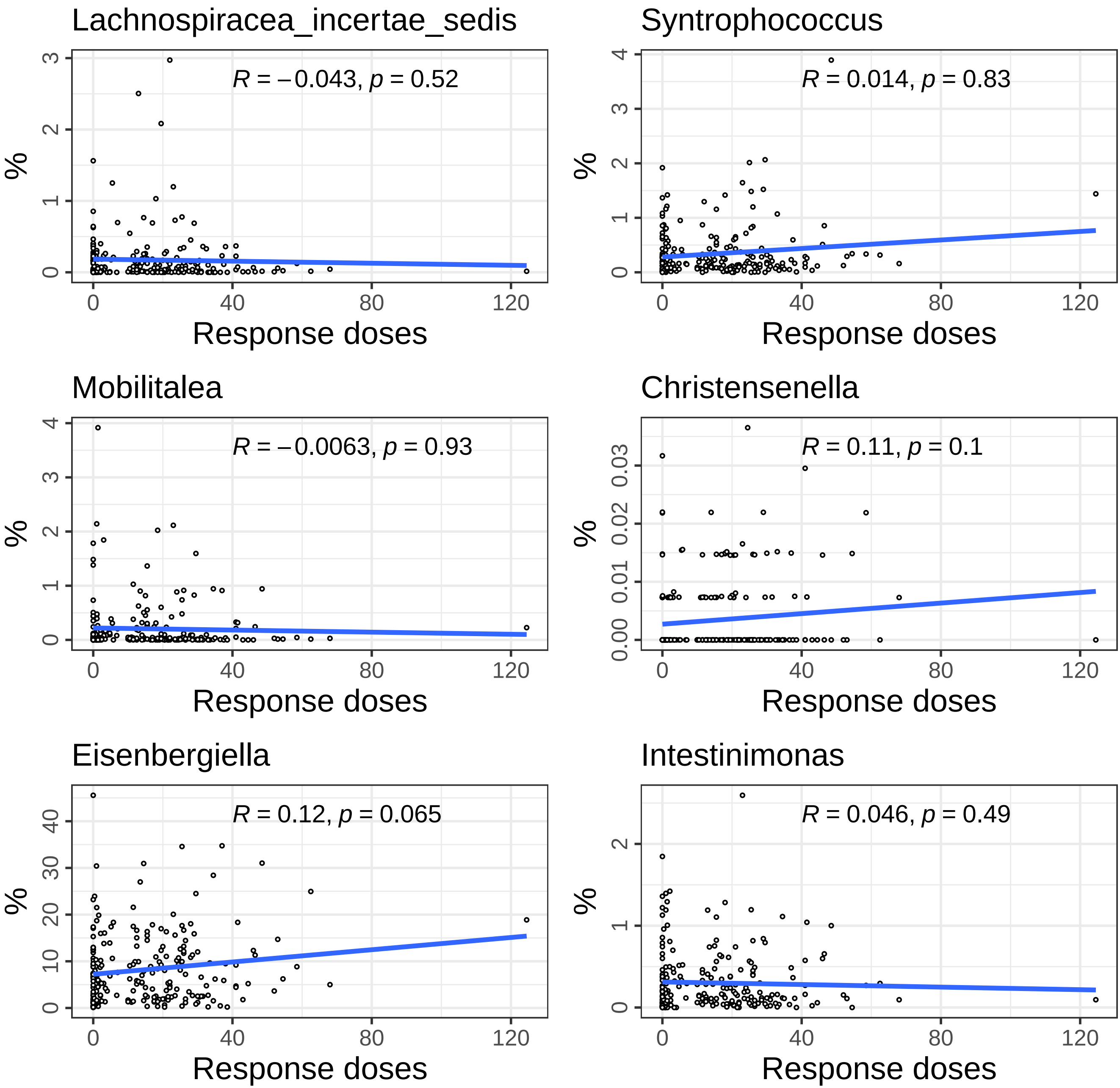
